# DNA mismatches reveal widespread conformational penalties in protein-DNA recognition

**DOI:** 10.1101/705558

**Authors:** Ariel Afek, Honglue Shi, Atul Rangadurai, Harshit Sahay, Hashim M. Al-Hashimi, Raluca Gordan

## Abstract

Transcription-factor (TF) proteins recognize specific genomic sequences, despite an overwhelming excess of non-specific DNA, to regulate complex gene expression programs^1–3^. While there have been significant advances in understanding how DNA sequence and shape contribute to recognition, some fundamental aspects of protein-DNA binding remain poorly understood^2,3^. Many DNA-binding proteins induce changes in the DNA structure outside the intrinsic B-DNA envelope. How the energetic cost associated with distorting DNA contributes to recognition has proven difficult to study and measure experimentally because the distorted DNA structures exist as low-abundance conformations in the naked B-DNA ensemble^4–10^. Here, we use a novel high-throughput assay called SaMBA (<u>Sa</u>turation <u>M</u>ismatch-<u>B</u>inding <u>A</u>ssay) to investigate the role of DNA conformational penalties in TF-DNA recognition. The approach introduces mismatched base-pairs (i.e. mispairs) within TF binding sites to pre-induce a variety of DNA structural distortions much larger than those induced by changes in Watson-Crick sequence. Strikingly, while most mismatches either weakened TF binding (~70%) or had negligible effects (~20%), approximately 10% of mismatches increased binding and at least one mismatch was found that increased the binding affinity for each of 21 examined TFs. Mismatches also converted sites from the non-specific affinity range into specific sites, and high-affinity sites into “super-sites” stronger than any known canonical binding site. These findings reveal a complex binding landscape that cannot be explained based on DNA sequence alone. Analysis of crystal structures together with NMR and molecular dynamics simulations revealed that many of the mismatches that increase binding induce distortions similar to those induced by TF binding, thus pre-paying some of the energetic cost to deform the DNA. Our work indicates that conformational penalties are a major determinant of protein-DNA recognition, and reveals mechanisms by which mismatches can recruit TFs and thus modulate replication and repair activities in the cell^11,12^.

## MAIN TEXT

A comprehensive survey of current crystal structures of protein-bound DNA (Methods) revealed that more than 40% of the complexes contain base-pair geometries that significantly deviate from the B-form envelope in naked DNA duplexes (**Fig. S1; Tables S1,S2**). The energy cost required to distort DNA must come from favorable intermolecular contacts that form upon complex formation^13,14^. This energetic cost could vary with sequence and contribute to protein-DNA binding affinity and selectivity^2,15,16^. Assessing these conformational penalties experimentally has proven difficult because it typically requires measuring the abundance of sparsely populated distorted conformations in the unbound DNA ensemble^4,5^, making them difficult to detect with conventional biophysical methods.

We developed an alternative, high-throughput approach to gain insights into the role of DNA conformational penalties in protein-DNA recognition. The approach—called SaMBA (<u>Sa</u>turation <u>M</u>ismatch <u>B</u>inding <u>A</u>ssay)—relies on using different types of mismatches to induce a variety of distortions to the DNA ensemble that are much greater than those attained by using different Watson-Crick sequences, and approach the distortions induced by proteins (**Fig. 1a,b,S2; Tables S3,S4**). For example, purine-purine mismatches widen the base-pairs, pyrimidine-pyrimidine mismatches constrict them, while wobble mismatches change the shear^17^. The distortions are further fine-tuned at the sequence level. For example, G(*syn*)-G(*anti*) mismatches form Hoogsteen-type base-pairs, A-A form distorted A(*anti*)-A(*anti*) base-pairs stabilized by a single hydrogen bond, while G-A can adopt a variety of configurations (**Fig S2**; Supplementary Discussion). Likewise, T-T mismatches form stable constricted wobbles whereas C-C form the least stable conformations. We reasoned that these different types of mismatches could increase the abundance of DNA conformational states that mimic the protein-bound conformation. By pre-paying the energetic cost for deformation, the mismatches could in turn increase the protein-DNA binding affinity, provided that the reduction in conformational penalty outweighs any destabilizing effects due to the potential loss of protein-DNA contacts. A conceptually similar strategy was recently used to assess conformational penalties in RNA-RNA association^10^. Since mismatches occur frequently in living cells^18,19^, SaMBA also provides a means by which to examine the impact of mismatches on protein-DNA landscapes and the biophysical mechanisms of these interactions, which have been recently suggested to play a role in mutagenesis^11,20^(**Fig. S3**).

**Figure 1.**
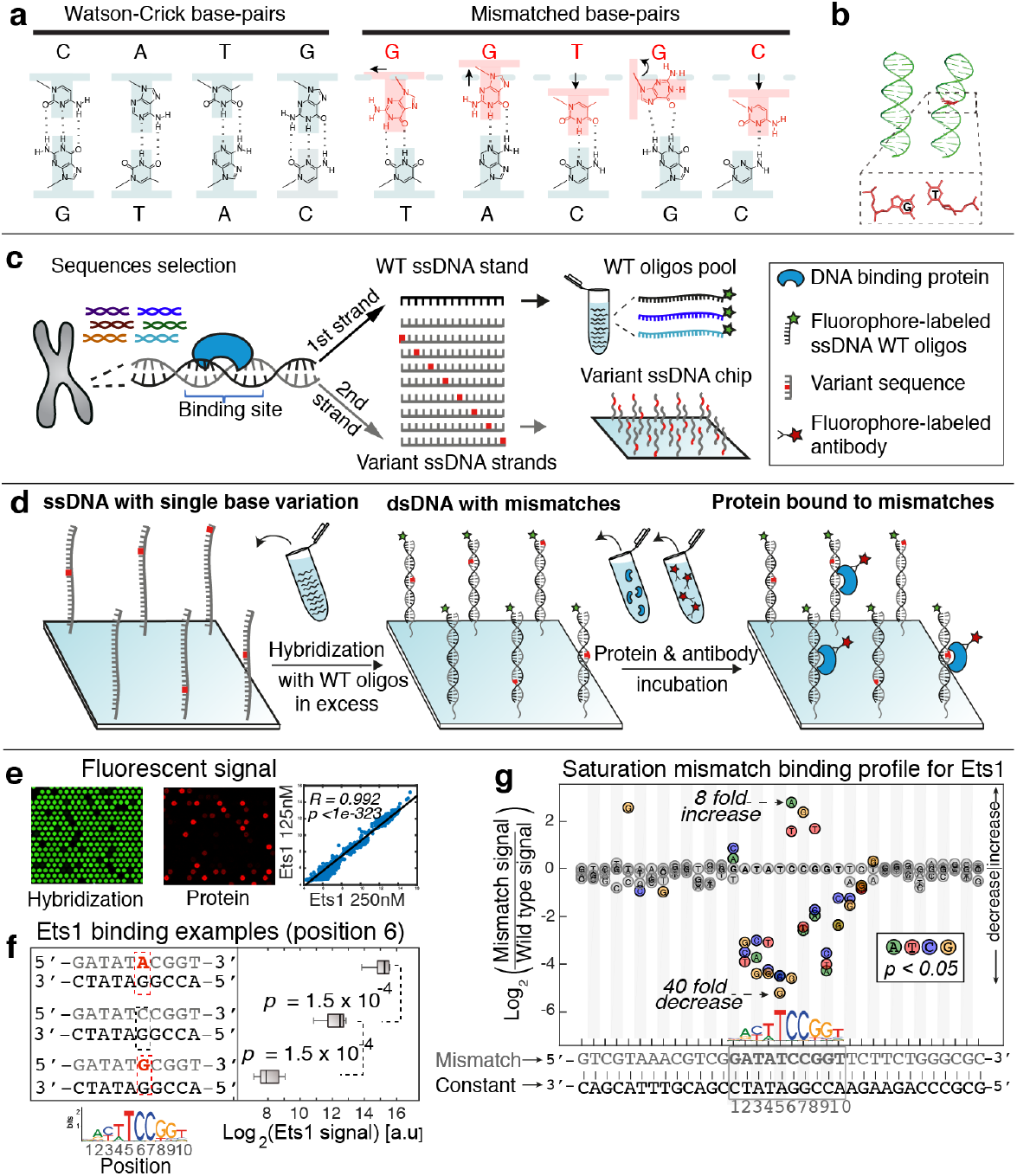
Overview of SaMBA, a high-throughput assay to measure the effects of mismatched base-pairs on protein-DNA binding. **(a)** Mismatched base-pairs (light red/blue) induce structural alterations such as base-pair shearing (G-T), helix diameter widening (G-A) or narrowing (T-C), *anti*-*syn* conformations (G-G) and paring destabilization (C-C) relative to Watson-Crick base-pairs (light blue). **(b)** Mismatches also influence the global structure of DNA by introducing kinks in the DNA helix. **(c)** In SaMBA, we first select DNA sequences containing protein-DNA binding sites of interest. 60-bp oligonucleotides containing each possible single-base variant, as well as the wild-type sequence, for one of the strands are synthesized on a DNA chip (Agilent) with ~15,000 DNA spots per chamber. In parallel, an oligo pool containing the wild-type complementary strands is produced, with a small fraction of oligonucleotides being fluorescently (Cy3) labeled. For the experiments described here, we chose DNA sites that are bound specifically by TF proteins (**Table S5; Data File 1**), according to universal protein-DNA binding microarray data21,22 (Methods). **(d)** The wild-type (WT) oligonucleotides, in excess, are hybridized to the chip, forming all possible single-base mismatched and WT duplexes. Next, the double-stranded DNA chip is incubated with the protein, which is detected with a fluorophore (Alex647)-conjugated antibody (similar to21), in order to enable quantitative detection of protein bound at each DNA spot. Previous work showed that the protein binding levels measured on DNA chips correlate well with independently measured binding affinities^21,22^. **(e)** Left: zoom-in image of a DNA chip showing the intensity of Cy3-labeled double-stranded DNA (green), illustrating uniform hybridization across all spots except for the dark control spots (see also **Fig. S4**). Middle: the Alexa647 signal (red) varies across DNA spots, illustrating different levels of protein binding. Right: scatterplot shows that SaMBA data is highly reproducible among replicate experiments. **(f)** The distributions of fluorescence signals for replicate spots are used to compare the TF binding at each mismatched sequence versus the corresponding wild-type sequence, using a Wilcoxon-Mann-Whitney test (see also **Fig. S5**). **(g)** Example of saturation mismatch binding profile showing the impact of single-nucleotide mismatches along one strand of a genomic Ets1 binding site on Ets1-DNA binding. Values are the log2 fold-change for the median intensity over 8 replicate spots. Positive values indicate an increase in Ets1 binding level compared to the WT sequence. Colored circles correspond to a significant change (Wilcoxon-Mann-Whitney test p-value < 0.05). Gray circles correspond to non-significant changes.

In SaMBA experiments, mismatches are generated by introducing all possible single-base variations (e.g. <u>G</u>-C to <u>A</u>-C, <u>T</u>-C, and <u>C</u>-C) in known DNA binding sites of TF proteins in a high throughput and unbiased manner on a high-density DNA chip (**Fig 1c-e,S4**; Methods). Protein binding measurements are then conducted directly on the chip (**Fig 1d-f**), and separate high throughput assays^21,22^ of protein binding to mutated Watson-Crick base-pairs are used to test for the impact of changing the DNA sequence.

We used SaMBA to obtain saturation mismatch binding profiles (**Fig. 1g,S5**) showing quantitative changes in protein-binding signal induced by the introduction of every possible mismatch to known TF binding sites and their flanking regions, for 21 TFs representing 14 distinct protein families (**Fig. 2a; Table S5; Data File 1**). The introduction of mismatches in sequences flanking TF binding sites had little to no effect on binding in most cases (84%), while mismatches within binding sites had wide-ranging effects. Most of the mismatches introduced within TF binding sites significantly weakened binding (67%); however, 8% of the mismatches increased binding, and at least one mismatch was found that increased the binding affinity for each of 21 examined TFs (**Fig. 2a,S6a)**. In the case of p53, where the increasing mismatch in SaMBA created a “super-site” stronger than the highest canonical p53 binding site^23^ (**Fig. S6a**), we confirmed the observed increase using fluorescence anisotropic binding affinity measurements (**Fig. S7**). Aggregate analysis at the mismatch-type level also revealed wide-ranging behaviors: every type of mismatch can increase or decrease binding, depending on the identity of the TF and the position of the mismatch within the binding site (**Fig. 2b**). The increased/decreased binding to mismatched DNA cannot be explained by sequence changes alone, since secondary rescue mutations that lead to mutated Watson-Crick base pairing have different, or even opposite effects on binding compared to mismatches (**Fig. 2c,S8**).

**Figure 2.**
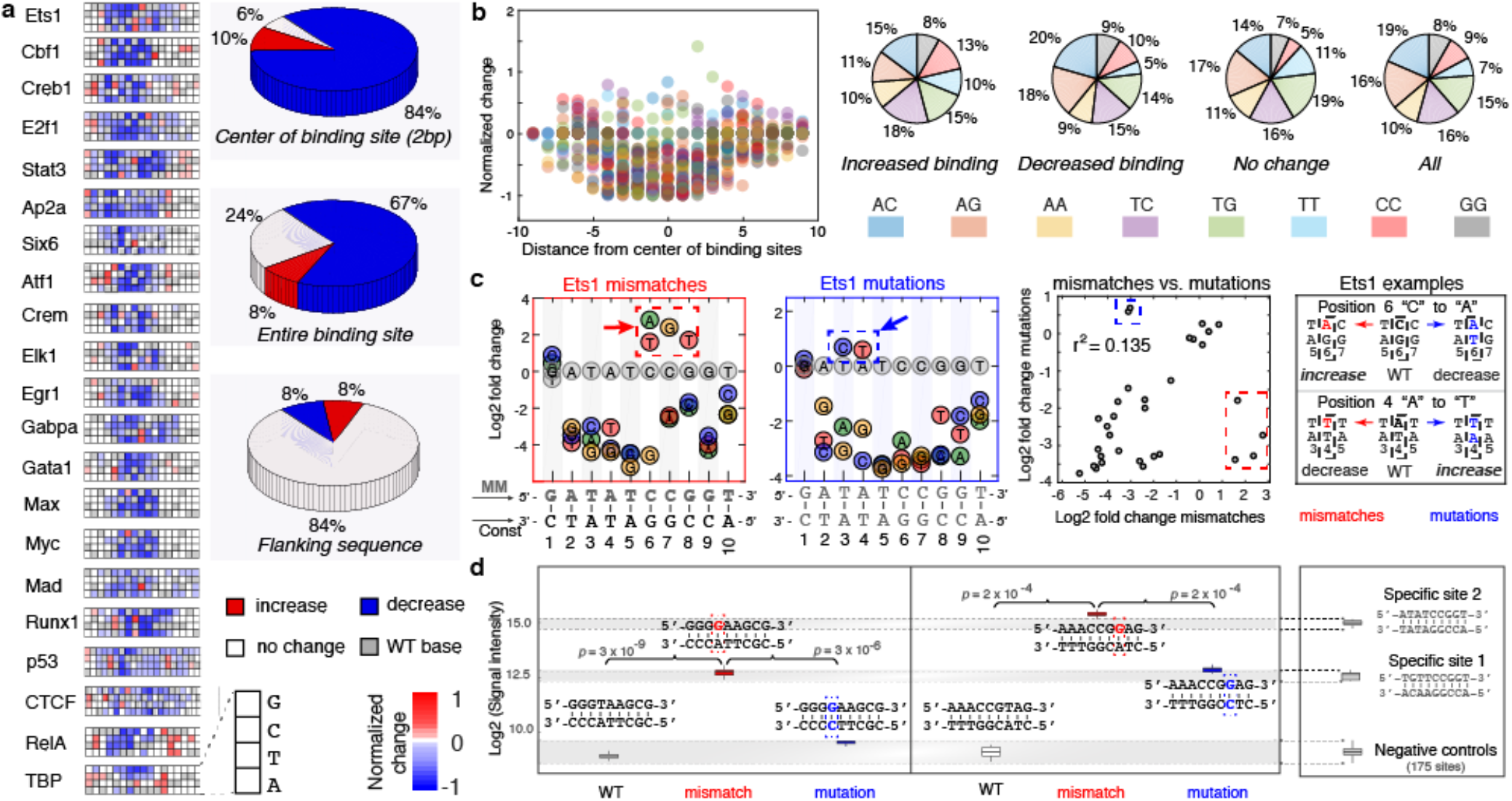
DNA mismatches have broad effects on TF binding, which cannot be explained by sequence changes alone. **(a)** Mismatch binding profiles for 21 TFs from 14 distinct protein families (**Table S5**). Heatmaps show the effects of mismatches on TF binding for short DNA sequences centered on TF binding sites. The color (red: increase, white: non-significant change, blue: decrease) in each entry represents the normalized log2 fold change in TF binding level due to a mismatch, compared to the wild-type sequence (Methods). The log2 fold changes (**Fig. S6a**) were scaled such that the biggest decrease in binding for each TF is set to −1 (dark blue). The wild-type bases are shown in gray. Pie charts show the distributions of significant increases (red), decreases (blue) and non-significant changes (white), within the center of the binding sites (top), the entire binding sites (middle), and in the sequences flanking the binding sites (bottom). **(b)** Normalized changes in TF binding due to mismatches, shown for all 21 TFs tested. Each color represents a different type of mismatch. Pie charts show the frequencies of the different mismatches across all 21 TF binding sites shown in the left plot. **(c)** Single base mismatches (left) and single base-pair mutations (second from the left) have distinct effects on Ets1 binding. The wild-type Ets1 site examined is shown underneath the plots. Scatter plot compares the effect of each mutation to the effect of its corresponding mismatch. Red rectangle marks significant increases unique to mismatches. Blue rectangle marks increases unique to mutations. Right: two examples illustrating the opposite effects of mismatches versus mutations. **(d)** Mismatches within DNA sequences bound with affinities in the non-specific range can increase Ets1 binding to generate strong binding sites that reach the level of known, specific Ets1 sites. We used universal protein-DNA binding microarray data22 to select a set of negative control sites that are non-specifically bound by Ets1, as well as the two positive control specific binding sites (**Fig. S6c**). Two example mismatches are shown (red), together with the corresponding secondary rescue mutations that lead to mutated Watson-Crick base pairs (blue). The Watson-Crick mutations have significantly smaller effects on Ets1 binding compared to the mismatches. In the first example, the binding level of the Watson-Crick mutation remains in the negative control range.

As a representative example, most mismatches within the Ets1 binding site shown in **Fig. 1g** significantly decreased protein binding, while those in the flanking regions generally had no effect. However, four of the 20 possible mismatches in the core Ets1 binding site (i.e. 20% of core mismatches) significantly increased, by as much as 8-fold, the protein binding level. For all four mismatches that increase Ets1 binding, the corresponding Watson-Crick rescue mutations decrease binding, relative to the parent sequence, by 7-fold or more (**Fig. 2c**). Similar results were robustly observed for other Ets1 sites (**Fig. S8b**). Thus, the changes in DNA binding affinity are mismatch-specific and cannot simply be attributed to changes in sequence. These results indicate that mismatch-induced structural deformations provide an additional layer of information about the role of DNA structure in protein-DNA recognition, beyond that which can be learned from analyzing the effects of sequence mutations in Watson-Crick DNA using traditional high-throughput methods^21,24^.

Due to the high density of the DNA chips used in our experiments, each SaMBA DNA library can accommodate binding sites for several TFs (**Fig. 1c**; **Data File 1**, Methods). Thus, each TF was tested not only against its specific binding site(s), but also a small number of non-specific sites, which were specific to other TFs (**Table S5; Data File 1**). For all proteins examined, the introduction of mismatches increased binding even at non-specific DNA sites and, surprisingly, in some cases the new binding levels were similar to those observed for specific binding sites (**Fig. 2d,S6b**). To verify the magnitude of such increases, for TF Ets1 we designed a DNA library that contained, in addition to Watson-Crick and mismatch-containing binding sites, a set of negative control probes lacking any site specifically bound by Ets1 (**Fig. S6c**). As shown in **Fig. 2d**, even when starting with Watson-Crick sites bound as weakly as the non-specific negative control group, the introduction of certain mismatches leads to strong protein binding, comparable to known Ets1 binding sites, thus effectively creating novel binding sites within non-specific DNA.

For a subset of 12 TF proteins with available crystal structures (**Table S6**), we examined whether the mismatch-induced perturbations that increase the binding affinity also induce distortions to the DNA structure that are similar to those induced by TF binding. We initially focused on p53, as p53-bound DNA structures have been extensively characterized^25,26^. The introduction of T-T and C-T to a high-affinity p53 binding site resulted in ~2-fold increase in the p53 binding level (**Fig. 3a,S6a**). Interestingly, these positions coincide with base-pairs shown to have a preference to adopt non-canonical Hoogsteen conformations^25,26^. Hoogsteen base-pairs represent an example of alternative lowly-populated conformations in apo-DNA ensemble, that form with abundance <1%^5,27^. The Hoogsteen pairing is achieved by flipping the purine base from an *anti* to *syn* conformation followed by a reduction of ~2Å in the helical diameter and C1′-C1′ distance (**Fig. 3a**). This reduction in the DNA diameter at the p53 binding site allows closer proximity of p53 monomers, thus stabilizing the p53 tetramer as compared to Watson-Crick pairs^26^.

**Figure 3.**
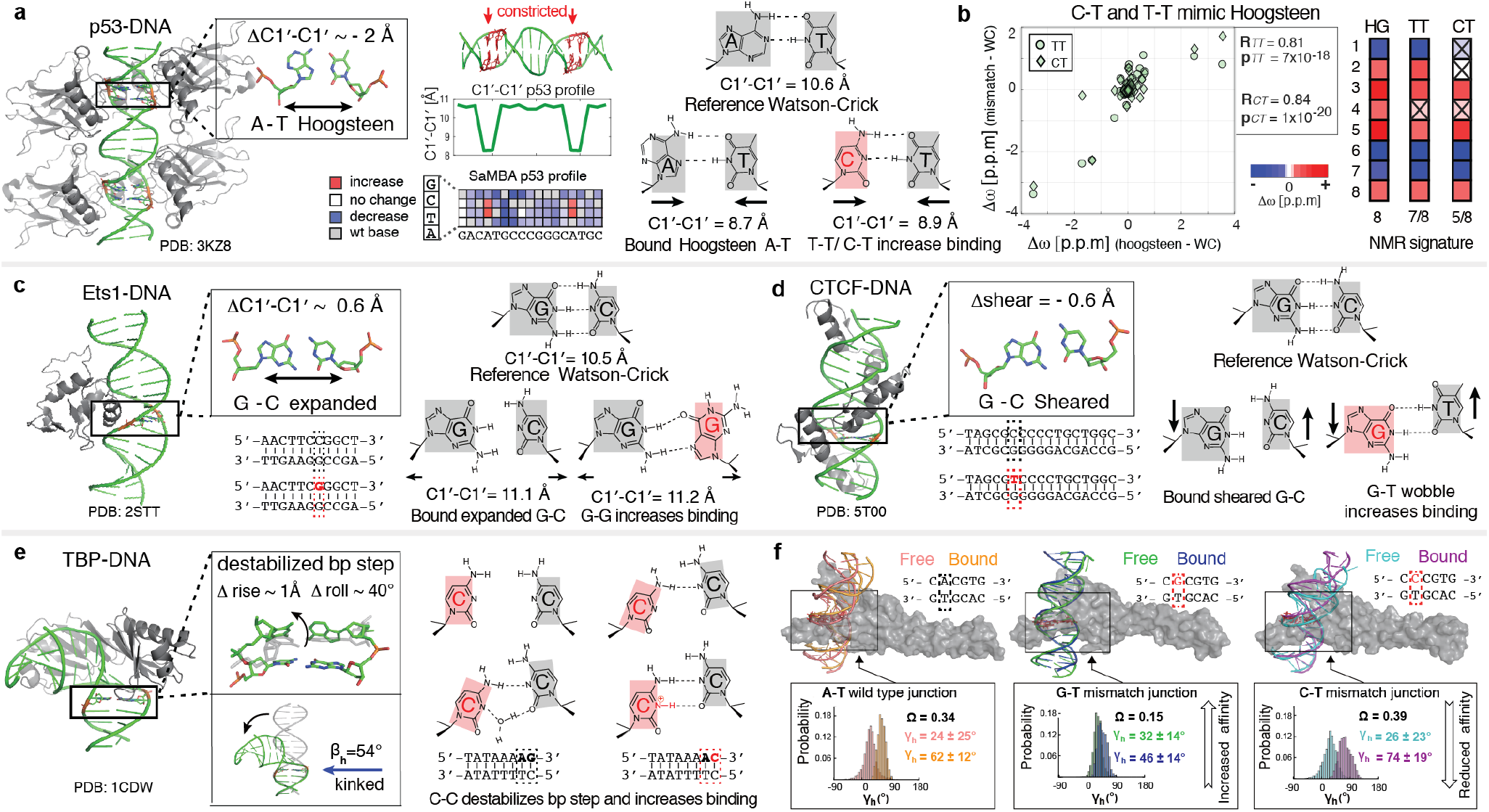
DNA mismatches that exhibit geometries similar to distorted base pairs in TF-bound DNA lead to increased binding affinity. **(a)** p53-DNA crystal structure shows a constricted Hoogsteen conformation at the A-T positions (black box, red arrows). C-T and T-T mismatches, which increase p53-DNA binding affinity, mimic Hoogsteen base pairing by constricting the C1′-C1′ distance. **(b)** NMR results confirm that T-T and C-T mismatches mimic Hoogsteen A-T geometry. Left: correlation plot of the chemical shift differences in the sugar C1′/C3′/C4′ carbons for T-T and C-T mismatches versus m1AT Hoogsteen base pairs, relative to the Watson-Crick base paired duplex. Right: heat map of the chemical shift signatures (Δω(m1AT-AT) > 0.5 ppm) (**Fig. S9a-d**) for base pair formation for m1A-T Hoogsteen, C-C, and T-T base pairs. Differences between m1A-T Hoogsteen and mismatched base pairs are marked with X. **(c)** Ets1-DNA crystal structure shows an expanded A-T base pair. The G-G mismatch at this position induces a similar deformation prior to Ets1 binding, thus increasing the Ets1 binding affinity. **(d)** CTCF-DNA crystal structure shows a deformed, sheared A-T base pair. The G-T wobble mismatch at this position induces similar shearing prior to CTCF binding, thus increasing the binding affinity. **(e)** TBP-DNA crystal structure shows destabilization at an ApG base pair step critical for TBP binding9,28,29. Replacing the G-C at this step with C-C, the mismatch with the lowest stacking propensity30, pre-destabilizes the DNA and increases TBP binding affinity. **(f)** DNA kinking plays an important role in Myc-DNA recognition. Compared to the WT site (left), a G-T mismatch leads to increased overlap between the kinking distributions for free and bound DNA, thus resulting in higher binding affinity (middle). In contrast, the C-T mismatch leads to a decreased overlap between the kinking distributions, thus decreasing binding affinity (right), consistent with experimental results. Histograms of the kinking direction (γ_h_) obtained from molecular dynamics simulations of free and bound DNA are shown along with their average values (Methods). The extent of overlap is analyzed using a revised Jensen-Shannon divergence score (Ω Methods). Representative structures of the DNA sites are shown for WT free (pink), WT bound (orange), G-T free (green), G-T bound (blue), C-T free (cyan), and C-T bound (magenta). The Myc/Max heterodimer is shown as a gray surface.

Remarkably, pyrimidine-pyrimidine mismatches also reduce the C1′-C1′ distance from ~10.5Å (for the Watson-Crick A-T pair) to ~8.5-9Å (for T-T and C-T) (**Fig. 3a**, **Table S3**). NMR analysis confirmed that the perturbations induced by A-T Hoogsteen base-pairs are indeed similar to those induced by T-T and C-T (**Fig. 3b,S9a-d**). These results indicate that by constricting the C1′-C1′ distance, T-T and C-T mismatches effectively mimic the p53-preferred Hoogsteen pairing in naked DNA, thus pre-paying some of the energetic penalty to form the preferred bound structure. These finding are in excellent agreement with a recent study in which modified bases were shown to induce Hoogsteen and increase p53 binding affinity in a similar manner^26^.

A more comprehensive analysis revealed mismatch-induced distortions similar to those induced by TF binding for approximately half of the 29 examined mismatches with increased binding (Methods; **Table S7**). For example, for Ets1, increased binding is observed when replacing an expanded C-G base-pair with a purine-purine G-G mismatch with increased C1′-C1′ distance (**Fig. 3c**); and for CTCF when replacing a sheared G-C base-pair with a corresponding G-T wobble (**Fig. 3d**). For TBP, prior studies showed that partial intercalation of Phe residues at two positions leads to loss of base-base stacking and formation of a sharp kink at two base-pair steps^9,28,29^. Remarkably, a C-C mismatch at one of these positions, introduced by the single-base variation of the SaMBA protocol, resulted in increased TBP binding (**Fig. 3e)**. This was expected given that C-C mismatches have the least favorable stacking interactions^30^. To evaluate the predictive power of these trends, we used not only single-base variations but also double-base variations to install C-C mismatches at every position within the same TBP binding sequence (5’-TATAAAAG-3’), and indeed we observed the expected increase in TBP binding for the other unstacked sites, while no increase in binding was observed at any of the remaining positions (**Fig. S9e-g**).

To examine mismatch-enhanced binding sites with no apparent structural mimicry to Watson-Crick bound DNA, we carried out molecular dynamics simulations on two example complexes for which G-T and C-T mismatches, which can be reliably modeled computationally (Methods), have opposite effects on TF binding. In the case of Myc (**Fig. 3f**), the G-T mismatch induces changes in both the free and TF-bound DNA ensembles, such that they are more similar compared to the parent Watson-Crick sequence, while C-T has the opposite trend. Thus, we cannot rule out that there are additional mechanisms for pre-paying conformational energy by changing both the free and bound DNA conformations. For Ets1, introducing a G-T mismatch leads to new protein-DNA contacts being formed during the simulation, suggesting a role for direct readout (**Fig. S10**). Although additional studies will be needed to separate these different contributions to the mismatch-enhanced binding observed by SaMBA, our results indicate that many binding increases will likely be explained by changes in DNA conformational penalties.

Our study provides strong evidence for DNA conformational penalties as an important and widespread determinant of protein-DNA binding affinity and selectivity. Our new assay can be extended to include distortions in DNA shape induced by multiple mismatches, insertions and deletions, damaged and epigenetically modified nucleotides, and thereby thoroughly investigate these penalties in a high-throughput and unbiased manner. In addition to their specific roles in the cell, DNA-bound proteins have been shown to act as roadblocks that can impede DNA repair and replication^11,12^. Therefore, high-affinity binding of TFs to DNA mismatches, oftentimes driven by reduced conformational penalties, may provide a biophysical mechanism for inhibiting the repair of specific mismatched sites, and consequently contributing to the formation of genetic mutations (**Fig. S3;** Supplementary Discussion). Thus, modulation of DNA conformational penalties could also play important roles in the mechanisms of genome stability.

## Supporting information

Methods and discussions

## Data and code availability

The data that support the findings in this study are available in the extended raw experimental data file, in Excel format. Any other relevant data are available from the corresponding author upon request.

Custom MATLAB code was use to analyze the data in this study, i.e. combine replicates and generate graphs. Custom Python code was used to analyze crystal structures, using X3DNA-DSSR. All code is available upon request.

## Acknowledgment

This work was funded by NIH grant R01-GM117106 (to R.G.), NIH grant R01-GM089846 (to H.M.A) and an Integrated DNA technologies postdoctoral fellowship (to A.A).

