## Supplementary material for "DNA mismatches reveal widespread conformational penalties in protein-DNA recognition": Suplemental figures

### Supplementary Figures for
### DNA mismatches reveal widespread conformational penalties in protein-DNA recognition

Ariel Afek[1,2], Honglue Shi[3], Atul Rangadurai[4], Harshit Sahay[1], Hashim M. Al-Hashimi[3,4]*, Raluca Gordan[1,2,5]*

[1]Center for Genomic and Computational Biology, [2]Department of Biostatistics and Bioinformatics, [3]Department of Chemistry, [4]Department of Biochemistry, [5]Department of Computer Science, Department of Molecular Genetics and Microbiology, Duke University. * Corresponding authors.

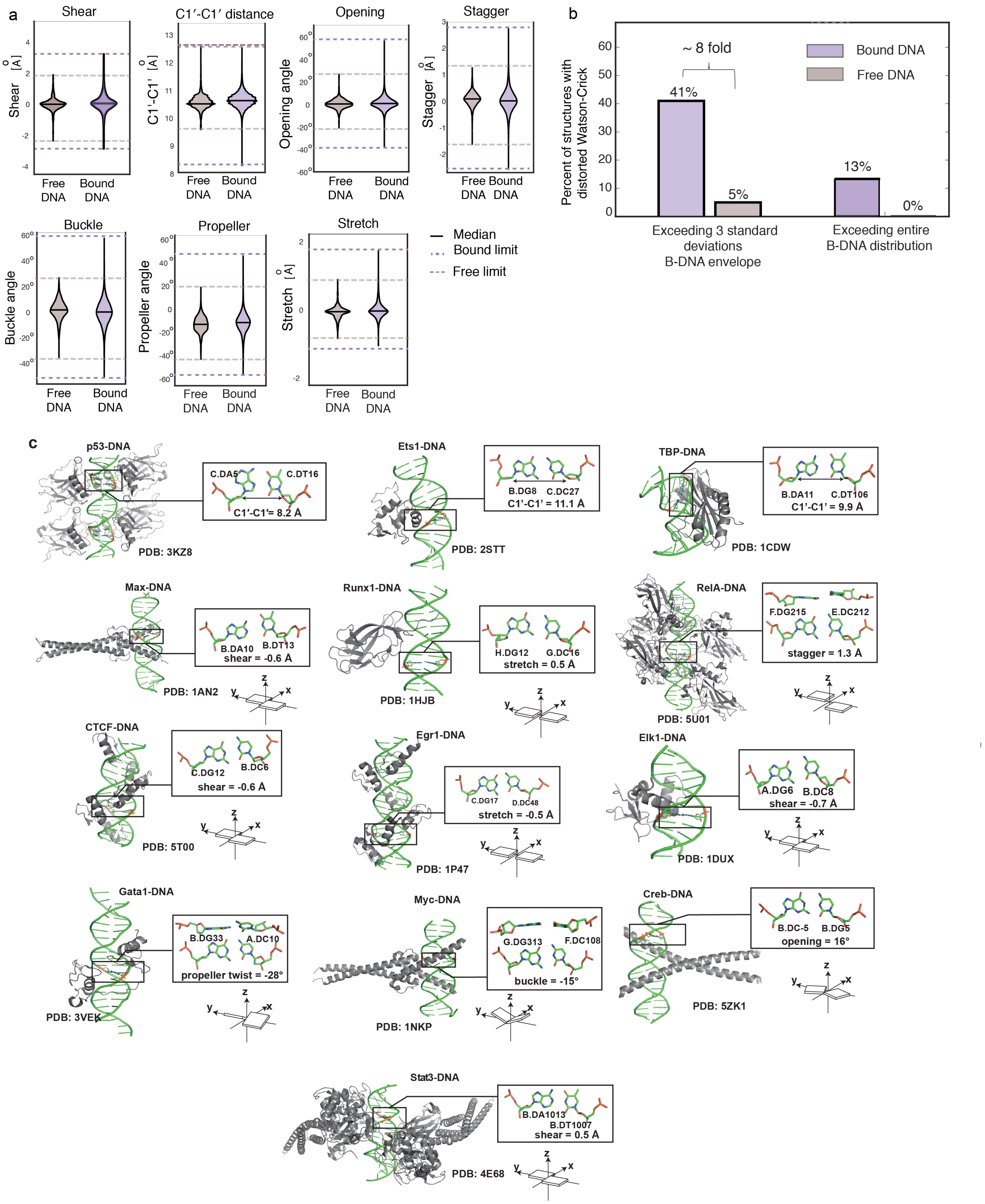

**Figure S1. (a) Distributions of base pair parameters in free and protein-bound DNA, from PDB[1] survey.** Solid lines denote the median value of the each parameter. Dashed lines denote the upper and lower bounds of the distribution for free (brown) and bound (purple) DNA. 1,736 protein-bound structures and 409 free B-DNA structures, all with resolution <3 Å, were used in the analysis.
**(b) Percentage of structures with base pairs outside the B-DNA envelope**. Among all 1,736 bound structures, 712 structures (41%) contain severe distortions of at least one base pair outside the free B-DNA envelope, with the envelope defined as at most 3 standard deviations above or below the mean (left purple bar). These distortions are ~8 times more common in bound structures compared to free DNA structures. (Using a less stringent definition of the B-DNA envelope, by considering 2 standard deviations above or below the mean, we found that 76% of the bound structures contain at least one base pair outside the free B-DNA envelope, approximately twice the frequency observed in free DNA.) Considering the full range of parameter values as defining the free B-DNA envelope, we identified 13% of the bound structures containing at least one base pair of extreme deformation, that were never observed in any free DNA structure (right purple bar).
**(c) Local deformations of base pairs observed in diverse transcription factor-DNA complex structures.** The complexes shown are: p53-DNA (PDB: 3KZ8), Ets1-DNA (PDB: 2STT), TBP-DNA (PDB: 1CDW), Max-DNA (PDB: 1AN2), Runx1-DNA (PDB: 1HJB), RelA-DNA (PDB: 5U01), CTCF-DNA (PDB: 5T00), Egr1-DNA (PDB: 1P47), Elk1-DNA (PDB: 1DUX), Gata1-DNA (PDB: 3VEK), Myc-DNA (PDB: 1NKP), Creb1-DNA (PDB: 5ZK1), and Stat3-DNA (PDB: 4E68). Left: 3D structures with the distorted base pairs highlighted in black boxes. Upper right: enlarged view of the base pair structures with their base pair parameters labeled. Lower right: schematic diagram of the corresponding base pair parameters.

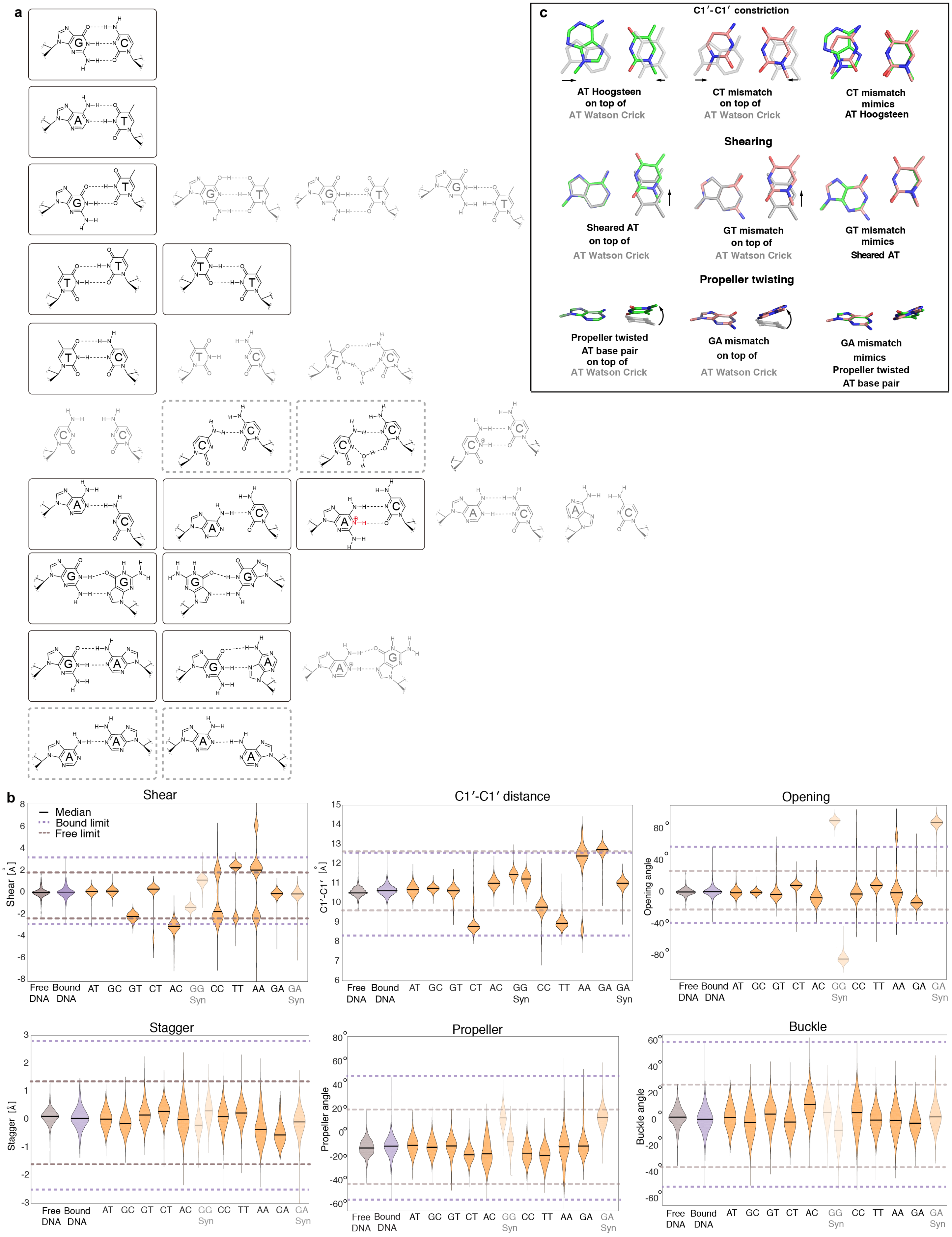
a
b
Shear
Median
Bound limit
Free limit
Shear [Å]
C1′-C1′ distance
C1′-C1′ [Å]
Opening
Opening angle
Stagger
Stagger [Å]
Propeller
Propeller angle
Buckle
Buckle angle
Free DNA
Bound DNA
AT
GC
GT
CT
AC
GG Syn
CC
TT
AA
GA
GA Syn
c
C1′-C1′ constriction
AT Hoogsteen on top of AT Watson Crick
CT mismatch on top of AT Watson Crick
CT mismatch mimics AT Hoogsteen
Shearing
Sheared AT on top of AT Watson Crick
GT mismatch on top of AT Watson Crick
GT mismatch mimics Sheared AT
Propeller twisting
Propeller twisted AT base pair on top of AT Watson Crick
GA mismatch on top of AT Watson Crick
GA mismatch mimics Propeller twisted AT base pair

**Figure S2. (a) Base pairing geometry of Watson-Crick base pairs and mismatches, obtained from a survey of crystal structures in the PDB[1].** Mismatches containing modified bases and those that were metal-mediated were excluded from the analysis. The predominant base pairing geometry under neutral pH conditions is indicated using a black box. Dashed boxes denote those mismatches whose base pairing geometries are uncertain. Minor base pairing geometries are shown in gray. See Methods for details. **(b) Distributions of base pair parameters in Watson-Crick and mismatched DNA, from MD simulations.** Solid lines denote the median value of the each parameter. Dashed lines denote the limits of structural parameter values for free and bound DNA structures from PDB (see Methods). Orange violins show the distributions of structural parameters for Watson-Crick and mismatched base pairs, based on MD simulations (see Methods). Observations from the MD simulations results: (1) The G-T mismatch remained wobble geometry with sheared conformation (|shear| around 2 Å) accompanied by a slight stretch during the MD simulation of both selected sequences (Methods). (2) The T-T mismatch shows wobble geometry with sheared conformation (|shear| around 2 Å). Different from G-T, the T-T mismatch shows rapid dynamic equilibrium of both wobble geometries with either one of the T shifted to the minor groove direction. Despite this rapid dynamic equilibrium, the T-T base pair is still constricted with C1′-C1′ distance 8-9.5 Å. (3) Similar to T-T, the C-T mismatch is also constricted with two H-bonds stably formed for most of the time. However, C-T mismatch can transiently adopt a high-energy conformation with only one H-bond formed and is not constricted anymore (C1′-C1′ distance ~10 Å) potentially due to the close contact between T-O2 and C-O2. The entire C-T MD trajectory is comprised of approximately 5% of these high-energy species. (4) The C-C mismatch is partially constricted with C1′-C1′ distance around 9.8 Å due to unstable H-bonding. (5) All pyrimidine-pyrimidine mismatches were still stacked in the helix without swing out of the helix in the MD trajectories. (6) The G-G mismatch does not experience *anti*-*syn* equilibrium during the simulation. The C1′-C1′ distance of G-G mismatch (G(*syn*)-G(*anti*) or G(*anti*)-G(*syn*)) is around 11.2-11.5 Å, which is larger than the canonical G-C base pair. (7) G(*anti*)-A(*syn*) is not constricted (C1′-C1′ distance around 11Å) and G(*anti*)-A(*anti*) reveals large C1′-C1′ distance around 12.8 Å.
**(c) Mismatches can mimic distorted base-pair geometries observed in protein-bound DNA.** Figure shows overlays of distorted (colored) and idealized Watson-Crick (gray) base pairs (left); mismatches (colored) and idealized Watson-Crick **(**gray**)** base pairs (middle); and mismatched and distorted Watson-Crick base pairs (right). The mismatched conformations presented are of free-DNA and were obtained from MD simulations (Methods). The C-T mismatch can mimic an A-T Hoogsteen base pair by constricting the C1′-C1′ distance (taken from PDB: 3KZ8). The G-T mismatch can mimic a sheared A-T base pair by shifting the T to the major groove direction (taken from PDB: 4MZR). The G-A mismatch can mimic a propeller twisted A-T base pair by twisting the A (taken from PDB: 4P0Q). In all cases, the Watson-Crick base pair is taken from an idealized B-form DNA structure constructed using 3DNA.

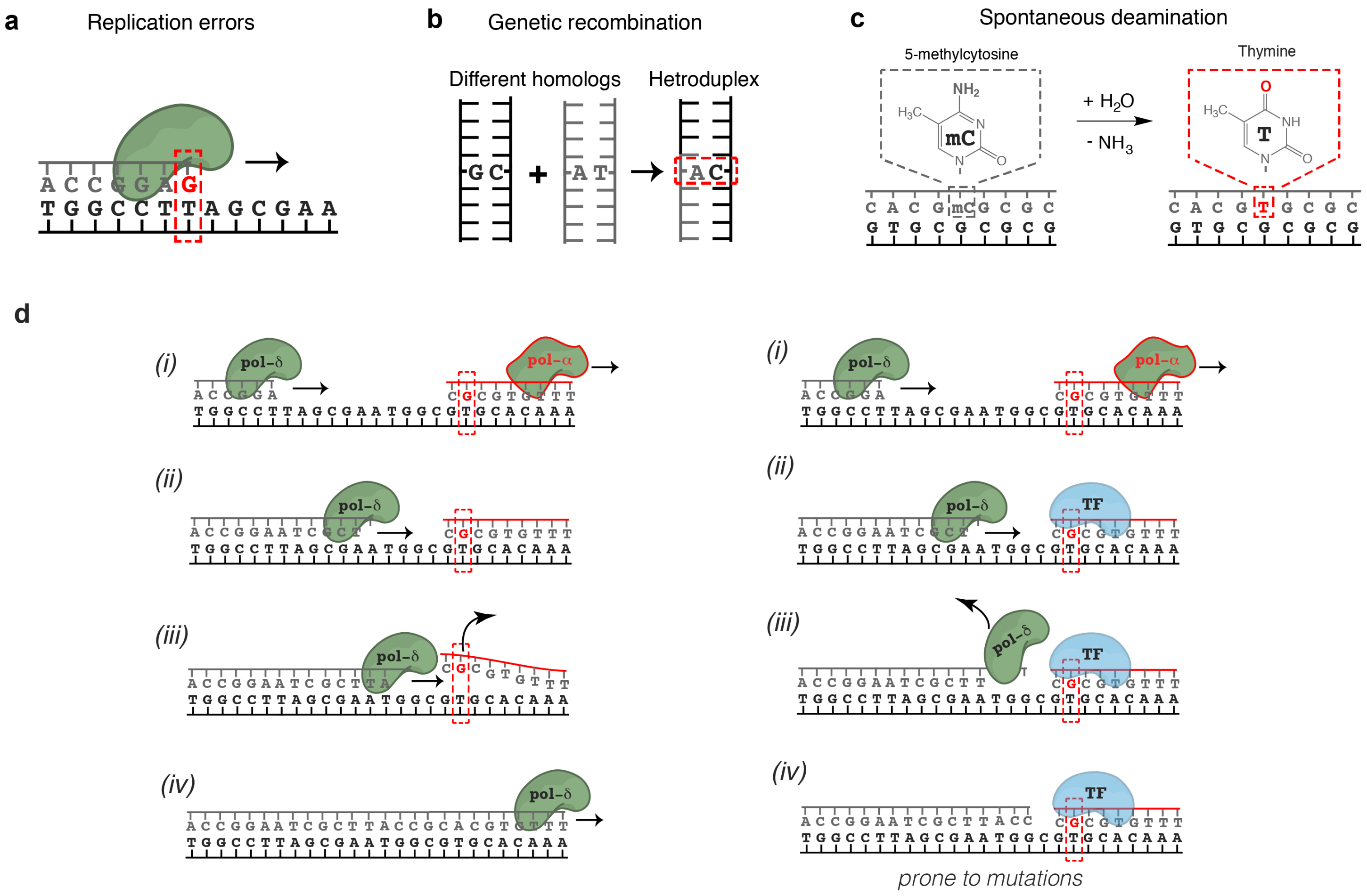


**Figure S3. DNA mismatches in the cell.**

**(a) Mismatches can result from misincorporation of bases during DNA replication by DNA polymerases.** The average rate at which replication errors are generated and escape proofreading is low in healthy cells (~$10^{-9}$), but high in certain cancers and cells with Pol-ε/Pol-δ mutations. Even in healthy cells, the rates of generation of individual mismatches vary by more than a million-fold[2] depending on the sequence context and the type of mismatch.

**(b) Mismatches result from genetic recombination.** A characteristic feature of homologous recombination is the exchange of DNA strands, which results in the formation of heteroduplex DNA. Mismatches can result from genetic recombination when the parental chromosomes contain non-identical sequences. In addition, mismatches can arise during DNA synthesis associated with recombination repair. The repair of these mismatches might be less efficient since it was previously shown[3] that there is a strong temporal coupling between DNA replication and mismatch repair but a lack of temporal coupling for heteroduplex rejection[3].

(**c**) **Spontaneous deamination** is common and estimated to occur 100-500 times per cell per day in humans[4]. G-T mismatches generated by deamination of 5-Methylcytosine (5-meC) are not repaired by the MMR pathway and have considerably lower repair efficiency[4]. The high rate of 5-meC deamination combined with their relatively slow repair in mammalian cells, contribute to making 5-meC a preferential target for point mutations (about 40-fold) compared to other nucleotides in the genome genome[5], and one of the major sources of the frequent C to T mutations observed in human cells[6].

**(d) Transcription factors bound to mismatched DNA could interfere with Pol-δ strand displacement activity.** Left: DNA synthesized by non-proofreading mismatch-prone Pol-α is normally displaced by the proofreading non-error-prone Pol-δ. Right: Reijns et al.[7] recently demonstrated that increased mutation signals arise from regions synthesized by Pol-α that contain TF binding sites. They suggested mismatched DNA synthesized by non-proofreading Pol-α is rapidly bound by TFs that act as barriers to Pol-δ displacement of Pol-α-synthesized DNA, resulting in locally increased mutation rates in subsequent rounds of replication.

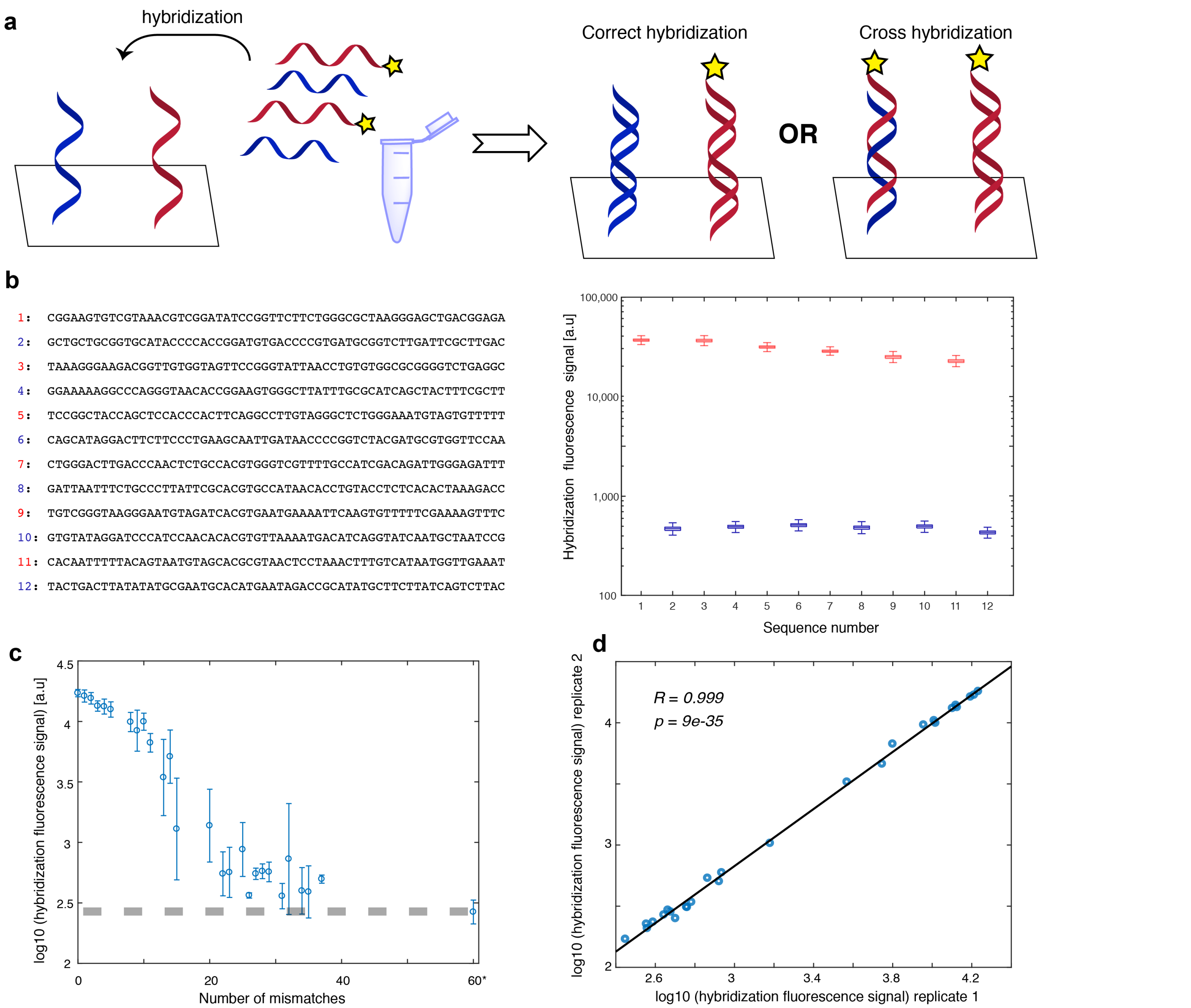


**Figure S4. Validation of DNA hybridization for wild-type and mismatched oligos during SaMBA.**
**(a) Schematic representation of our experimental workflow to detect cross-hybridization.** To check whether significant cross-hybridization occurs (i.e. whether certain oligonucleotides hybridize with non-target complementary oligonucleotides), we designed an experiment in which only certain oligonucleotides (red) were fluorescently labeled, and the others (blue) were not. If significant cross-hybridization occurred, we would have detected fluorescent signal on chip even for sequences without fluorescent-complements in the hybridization solution (i.e. for the sequences shown in blue).
**(b) No significant cross-hybridization was experimentally detected**. Left: list of 12 sequences used in the hybridization solution of one SaMBA experiment; red numbers are used for the fluorescently labeled oligonucleotides, and blue for the unlabeled sequences. Right: the measured fluorescent signal levels from the hybridization of these 12 sequences and their complementary sequences on the chip. For the sequences on the chip for which their complement is non-fluorescently labeled, the fluorescent signal is practically undetectable (blue), and it is several order of magnitude lower than the sequences with a labeled complementary strand (red). This demonstrates that no detectable cross-hybridization occurs.
**(c) The effect of mismatches on hybridization.** To estimate the efficiency of our current hybridization protocol, we measured the hybridization signal of one specific sequence in the solution (sequence #3 for library “v1”) , to different sequences containing multiple mismatches (0 to ~40), and a completely different sequence (‘60*’). As expected, the hybridization is less efficient for sequences with large numbers of mismatches. However, for small numbers of mismatches the hybridization is highly efficient. Longer incubation time, higher oligonucleotide concentration, and normalization of the signal could enable the usage of SaMBA for measuring TF binding to larger numbers of mismatches. Each data point in the plot shows medians and standard deviations over multiple sequences containing the same number of mismatches, with each sequence present in 6 replicate spots. The mismatches were introduced in a stochastic procedure. Briefly, we introduced N random base changes (N=1,2,3,4,5,10,15,25,35,45) to sequence #3, and we repeated this procedure ten times for each N to generate a total of 10 different sequences for each N. This procedure produced random sequences with the number of random mismatches in each duplex ranging between 1 and 37 compared to the original wild-type sequence.
**(d) Hybridization signal is highly reproducible**. We repeated the hybridization procedure described in (c) a second time; the correlation of hybridization signals between the two replicate experiments was very high ($R^2$=0.99). Plot shows median values based on data shown in panel (c).

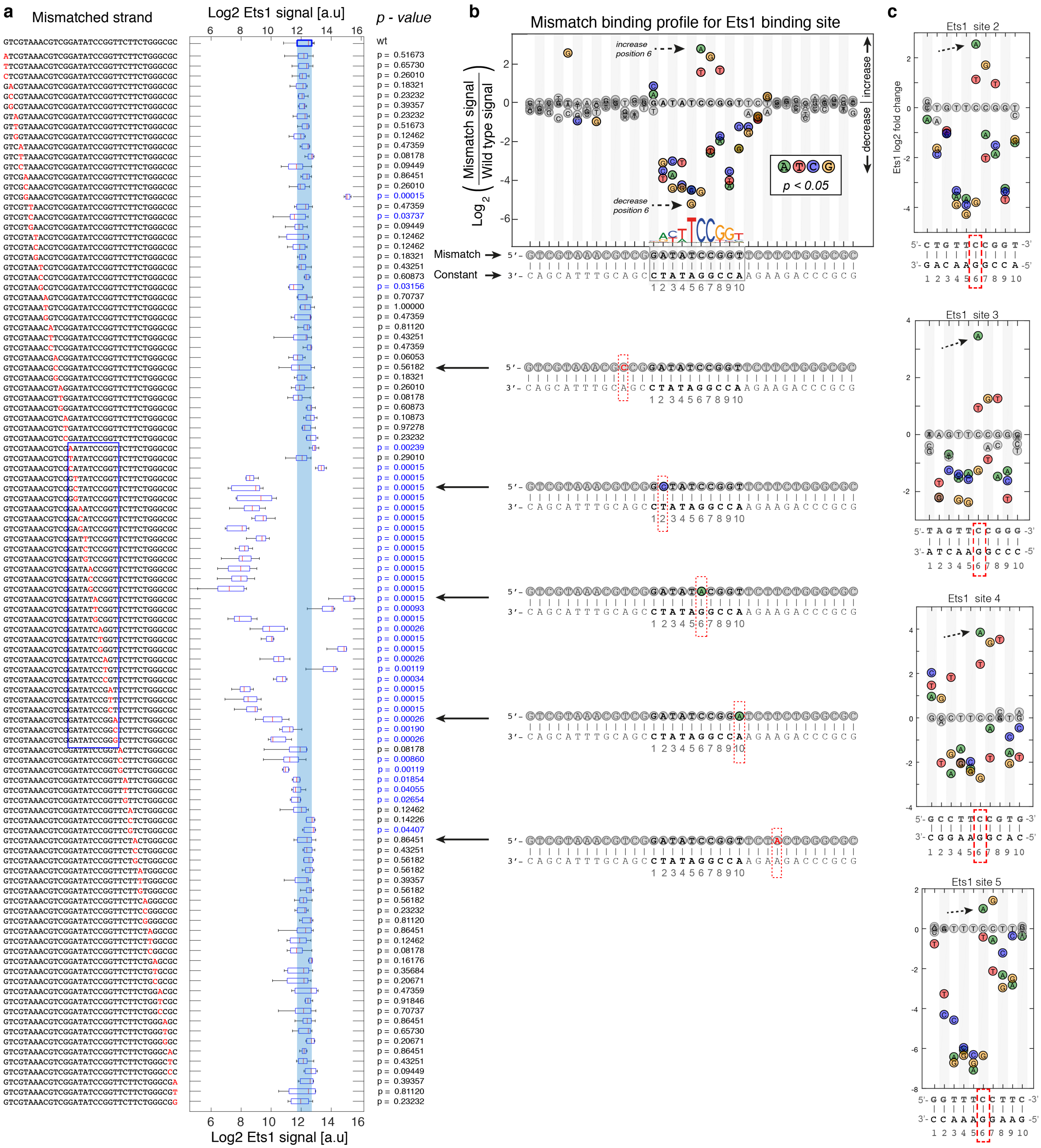


**Figure S5. (a,b) Detailed SaMBA profile for a TF binding site.**

(a) For each TF binding site of interest (here, an Ets1 binding site), the SaMBA DNA library synthesized on the chip contains both the wild-type site (i.e. the perfect complement of the strand used for hybridization; top left) and each possible 1-nt variant (red) in 8-20 replicates. A box plot presenting the Ets1 binding distribution for each sequence is shown. The central mark indicates the median, and the bottom and top edges of the box indicate the 25th and 75th percentiles, respectively. The whiskers extend to the most extreme data points not considered outliers. Vertical light blue line highlights the distribution of TF binding signals observed for the wild-type (WT) site. P-values on the right of the boxplots represent the two-sided Wilcoxon-Mann-Whitney test p-values for the differences between TF binding signal distributions for the WT versus the particular mismatch. Examples are presented on the right of the plot.

(**b**) A more compact representation with similar information as in (a). The height of each circle represents the log2 ratio between the median value of Ets1 binding to the WT site versus each mismatch. The letter inside each circle represents the variation introduced at that position. Mismatches with significant p-value ($p<0.05$) are shown in color.

**(c) SaMBA profiles for additional Ets1 binding sites.** We measured the effect of mismatches for four additional Ets1 binding sites We can see that although the profiles for different cores are quantitatively different and dependent on the flanks, the trends for increased binding due to mismatches are similar. For example, for all cases the A-G mismatch at position 6 significantly increases Ets1 binding.

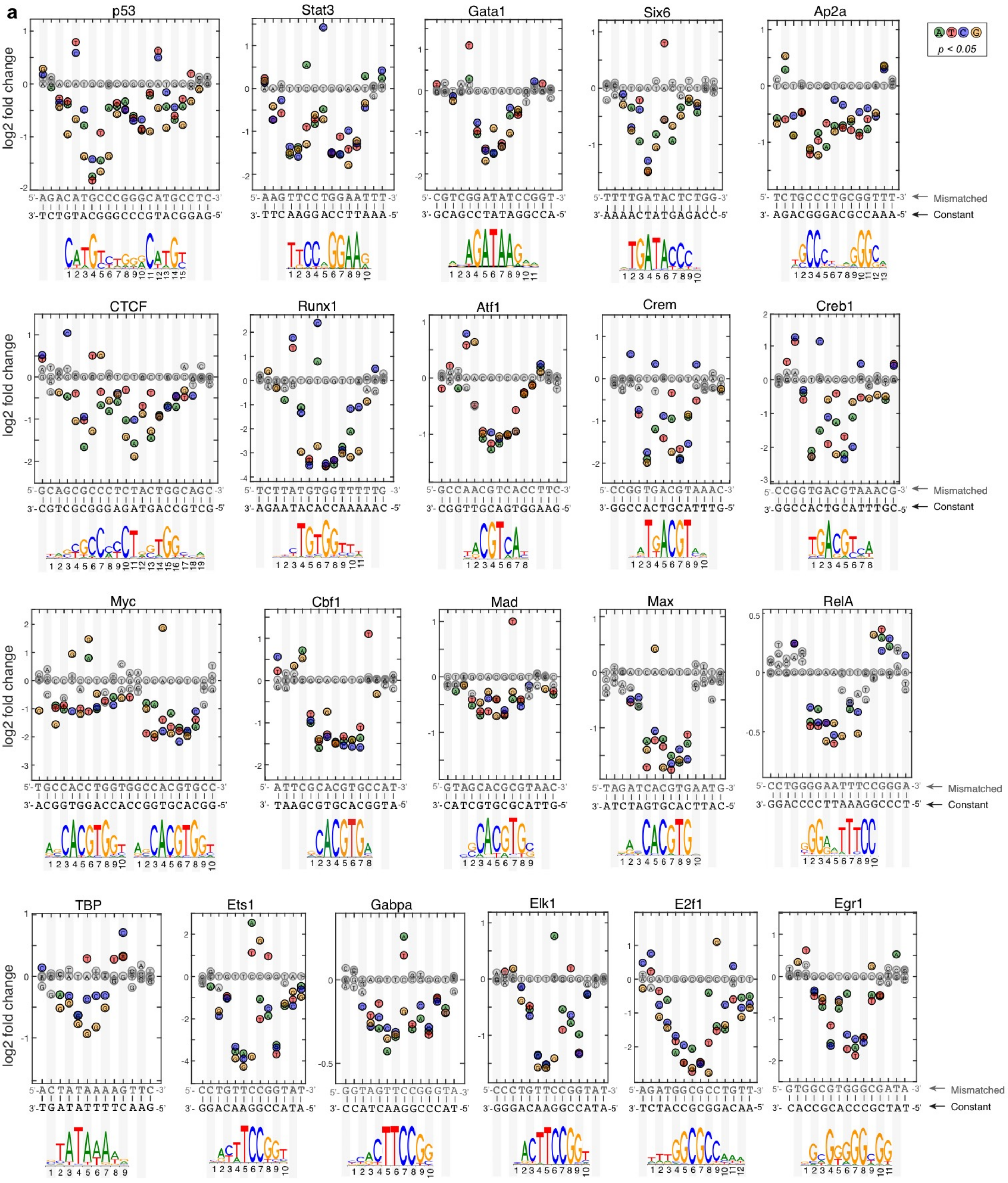

a
p53
Stat3
Gata1
Six6
Ap2a
CTCF
Runx1
Atf1
Crem
Creb1
Myc
Cbf1
Mad
Max
RelA
TBP
Ets1
Gabpa
Elk1
E2f1
Egr1
log2 fold change
Mismatched
Constant
p < 0.05

**b**

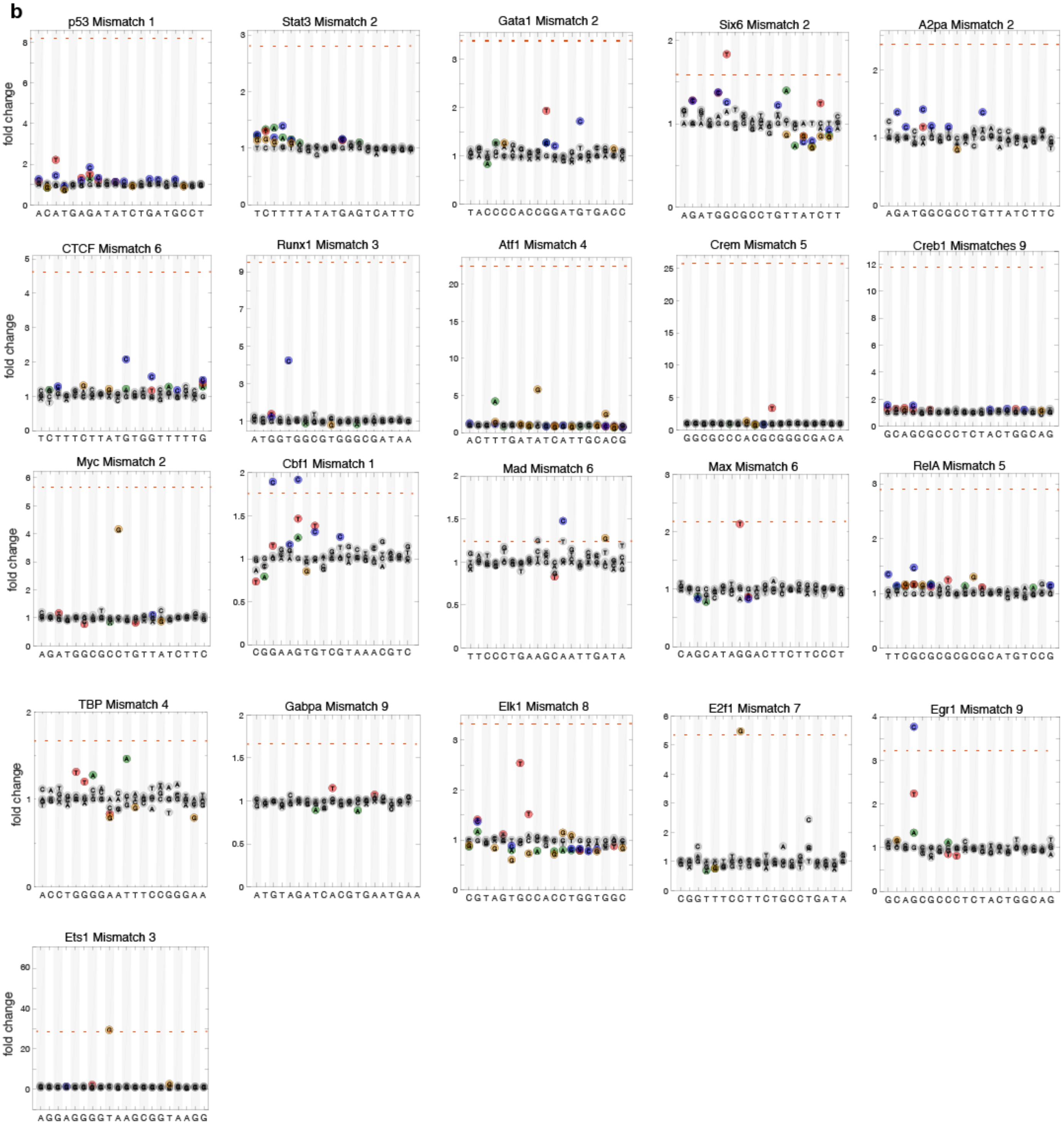
p53 Mismatch 1
Stat3 Mismatch 2
Gata1 Mismatch 2
Six6 Mismatch 2
A2pa Mismatch 2
CTCF Mismatch 6
Runx1 Mismatch 3
Atf1 Mismatch 4
Crem Mismatch 5
Creb1 Mismatches 9
Myc Mismatch 2
Cbf1 Mismatch 1
Mad Mismatch 6
Max Mismatch 6
RelA Mismatch 5
TBP Mismatch 4
Gabpa Mismatch 9
Elk1 Mismatch 8
E2f1 Mismatch 7
Egr1 Mismatch 9
Ets1 Mismatch 3
fold change

**c**

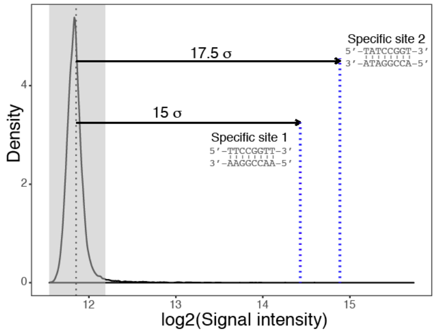
17.5 σ
Specific site 2
5'-TATCCGGT-3'
3'-ATAGGCCA-5'
15 σ
Specific site 1
5'-TTCCGGTT-3'
3'-AAGGCCAA-5'
Density
log2(Signal intensity)

**Figure S6. SaMBA profiles reveal increased binding due to mismatches for all 21 TFs tested**.

**(a)** Profiles showing the impact of single-nucleotide mismatches along one strand of genomic binding sites on the binding of 21 TFs. Values are the log2 fold-change for the median intensity (over 8-20 replicates, depending on the SaMBA library; see Methods). Positive values indicate an increase in binding compare to the WT sequence. Colored circles correspond to a significant change (Wilcoxon-Mann-Whitney test p-value < 0.05). Gray circles correspond to a non-significant change. Sequence changes to generate mismatches were made on one strand labeled as 'Mismatched' on the right, which is colored gray. The strand that was unchanged is shown in black and labeled as 'Constant'. The binding motif logo according to the Jaspar database[8] is shown for each TF below its SaMBA profile.

**(b)** SaMBA profiles for mismatches in non-specific binding sites. The mismatch strand is shown underneath each profile.

**(c)** Distribution of Ets1 binding signals for all possible 8-bp DNA sequences, as measured using an universal PBM assay [9,10].. Values shown are log2 of the median fluorescent intensities for each 8-mer. The shaded area represents the mean of the distribution +/- 2 standard deviations, where the binding can be assumed to be non-specific; all 8-mers in this range have a universal PBM enrichment score (E-score) < 0.3[10]. The two blue dashed lines represent two specific Ets1 sites that were also chosen, which fall within the top 0.1% and the top 0.01% of all 8-mers, respectively binding intensities of the two selected specific sites are at least 15 standard deviations above the average Ets1 binding signal. These two sites, as well as 175 random sites from the shaded non-specific region were selected as positive and negative controls, respectively, for an Ets1 SaMBA experiment to verify the magnitude of the binding increases due to mismatches in non-specific DNA.

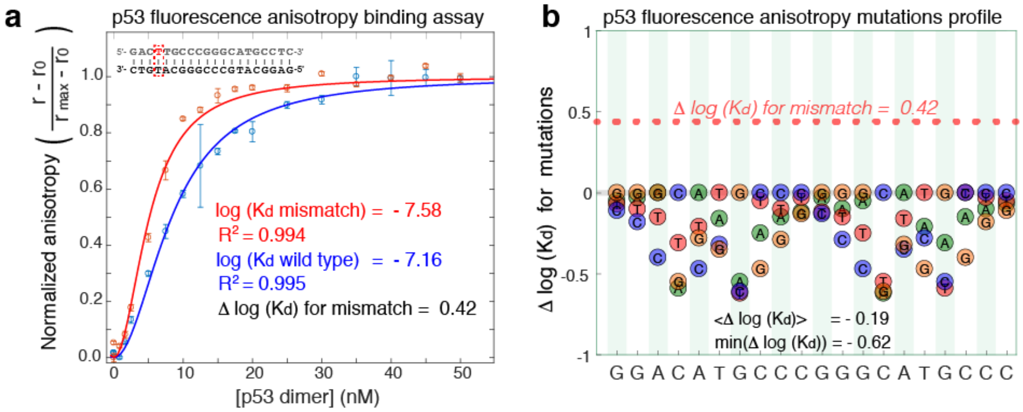


**Figure S7. Fluorescence anisotropy measurements for p53.**

**(a)** The binding of p53 protein to wild-type DNA binding site duplex (21-bp) with fluorescein dye attached to the 5′-terminus, and to an identical duplex with a T-T mismatch, also with 5'-fluorescein dye, was measured using fluorescence anisotropy. Kd values were obtained by fitting the normalized anisotropy across 17 different p53 dimer-concentration to the Hill equation $\theta = \frac{[P]^n}{Kd+[P]^n}$ , with Hill coefficient of n=2, as in [11].

**(b)** Δlog(Kd) for different mutations of p53 WT binding site. The data used for this figure was taken from [12] (Table 2). The fluorescence anisotropy measurements performed in [12] showed that the average Δlog(Kd) for mutations is ~0.2, and the maximum increase due to mutation is Δlog(Kd) of ~0.6. The observed increase we see due to mismatch (Δlog(Kd) of ~0.4, red asterisks; which is equivalent to a change of 0.97 kT in the binding energy), has a similar magnitude to the observed mutations, but the opposite direction.

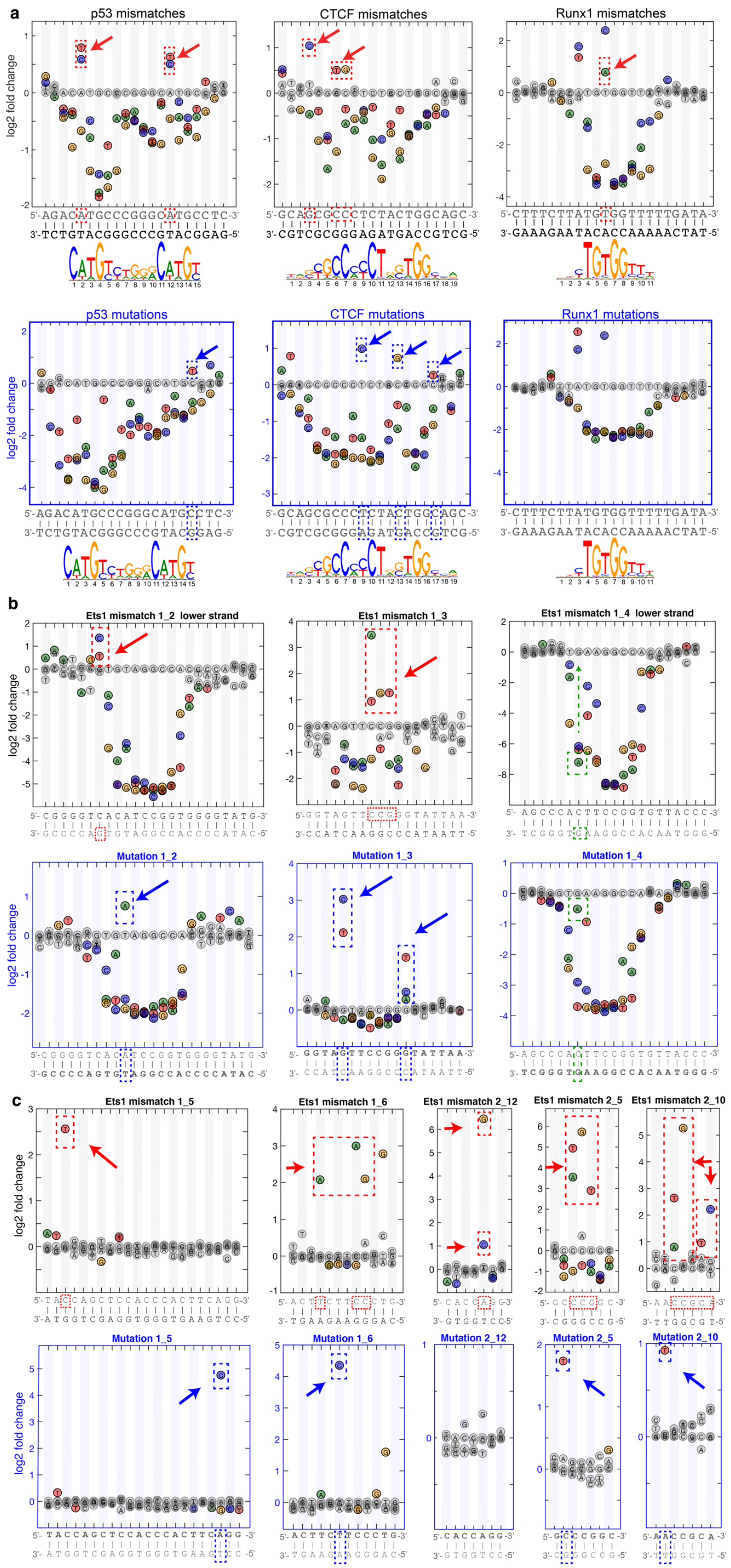


**Figure S8. Comparisons of SaMBA mismatch profiles and saturation mutagenesis profiles for p53, CTCF and Runx1, as well as Ets1 in additional specific and non-specific sites.**

**(a)** Mismatch and mutation profiles for p53, CTCF and Runx1**.** Red rectangles highlight increased TF binding caused only by mismatches, while blue rectangles highlight increased binding unique to mutations.

**(b)** The effect of mismatches and the equivalent mutations for three additional Ets1 binding sites (on top of the one presented in the main text). Once again we see that the increases in binding due to mismatches (red rectangle) versus mutations (blue rectangle) occur at different positions. We can also see cases where both the mismatch and the mutation cause a decrease in binding, but the magnitude of the decrease is very different (top right, green rectangle).

**(c)** Mismatches and mutations have different effects on Ets1 binding at non-specific sites. We can see examples for sites in which no mutation in the binding site has a strong effect, but the corresponding mismatches can increase TF binding by more than 60-fold (example 2_12).

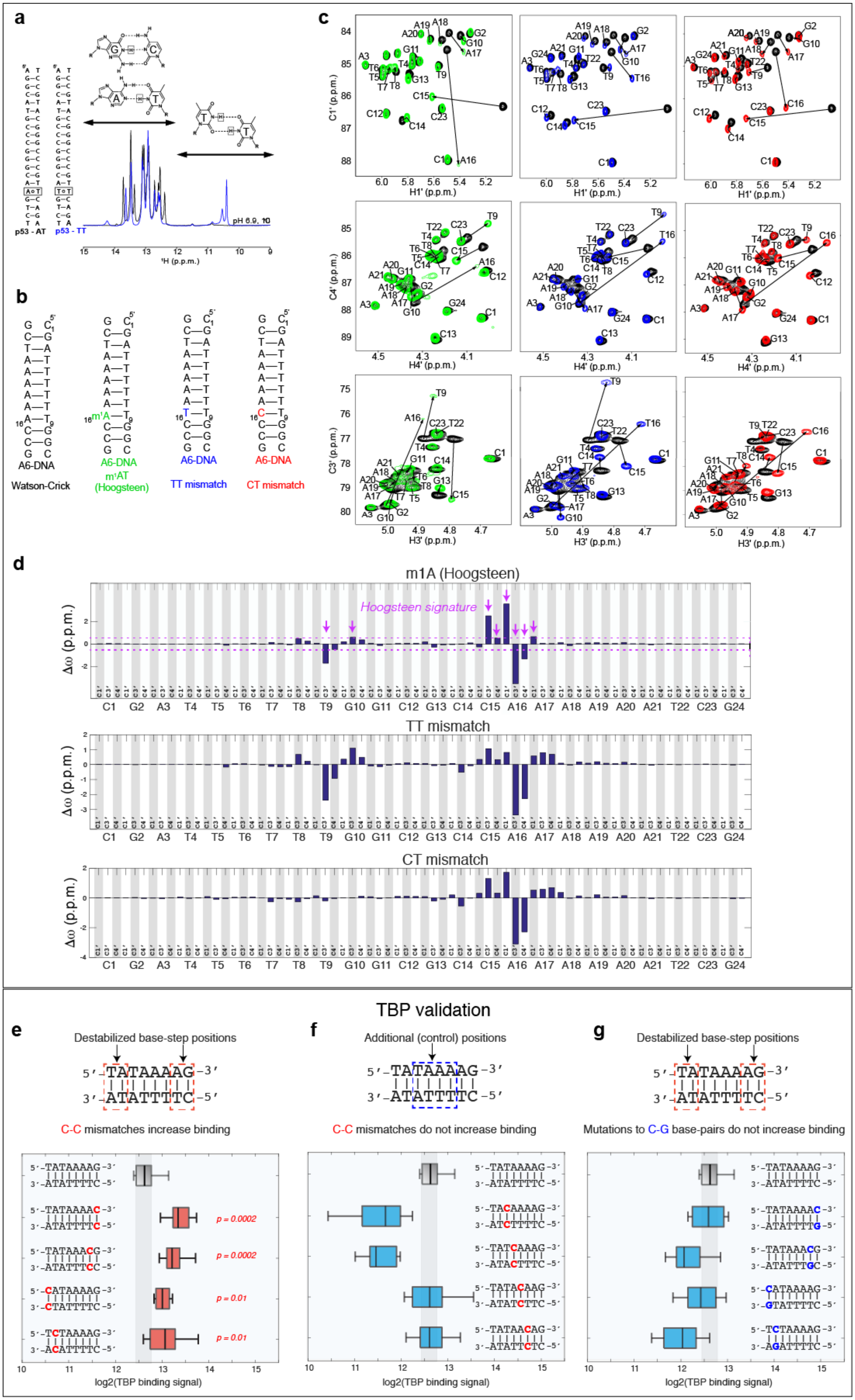

a
p53 - AT
p53 - TT
pH 6.9, 10
¹H (p.p.m.)
b
A6-DNA
Watson-Crick
A6-DNA
m¹AT
(Hoogsteen)
A6-DNA
TT mismatch
A6-DNA
CT mismatch
c
H1' (p.p.m.)
C1' (p.p.m.)
H4' (p.p.m.)
C4' (p.p.m.)
H3' (p.p.m.)
C3' (p.p.m.)
d
m1A (Hoogsteen)
Hoogsteen signature
Δω (p.p.m.)
TT mismatch
CT mismatch
C1 G2 A3 T4 T5 T6 T7 T8 T9 G10 G11 C12 G13 C14 C15 A16 A17 A18 A19 A20 A21 T22 C23 G24
TBP validation
e
Destabilized base-step positions
5'-TATAAAAG-3'
3'-ATATTTTC-5'
C-C mismatches increase binding
p = 0.0002
p = 0.0002
p = 0.01
p = 0.01
log2(TBP binding signal)
f
Additional (control) positions
5'-TATAAAAG-3'
3'-ATATTTTC-5'
C-C mismatches do not increase binding
log2(TBP binding signal)
g
Destabilized base-step positions
5'-TATAAAAG-3'
3'-ATATTTTC-5'
Mutations to C-G base-pairs do not increase binding
log2(TBP binding signal)

**Figure S9. Validation of the proposed hypothesis that mismatches can enhance TF binding by mimicking bound DNA distortions.**
**(a-d) NMR validation showing that T-T and C-T mimic the reduced C1'-C1' distance observed in p53-bound DNA[13,14].**
(a) NMR spectra for wild type and T-T mismatch-containing unbound p53 site. In the above spectra, each signal arises from the hydrogen of an N-H group on the DNA bases that are paired, typically from non-terminal Guanines or Thymines. Differences in the positions of the peak reflect different chemical environments of each N-H proton. The signals between 12 and 15 ppm typically arise from Watson- Crick base pairs in which the hydrogen bond acceptor for the N-H proton is another nitrogen atom on the complementary base. The signals between 9 and 12 ppm typically arise from N-H protons that are hydrogen bonded to oxygen atoms such as in mis-paired bases. For the T-T (red) samples we see two peaks corresponding to the N-H imino protons on the two Thymines 10-12 ppm, while such peaks are not visible for the A-T samples. The fact that we do see the signal between 9-12 ppm for T-T suggests that the N-H proton is directly bonded to another O atom on the partner base. This can only happen when the C1′-C1′ distance decreases relative to Watson-Crick base pairs (as two pyrimidine bases are smaller than a purine and a pyrimidine) to form the wobble base-pairing geometry.
(b) Secondary structures of A6-DNA variants used for chemical shift measurements.
(c) Chemical shift overlays of the 2D HSQC NMR spectra of the C1', C4' and C3' regions for A6-DNA m1A (left, green), A6-DNA TT (middle, blue) and A6-DNA CT (right, red) with unmodified A6-DNA (black) at pH 6.9, 25 °C.
(d) Bar plots of the chemical shift differences of the C1'/C3'/C4' carbons between A6DNA m1A (top), A6-DNA TT (middle) and A6-DNA CT (bottom) and A6-DNA. Purple lines indicate the chemical shift cutoff (+/-0.5ppm) used for defining chemical shift signatures for Hoogsteen base pair formation.
**(e-g) Independent validation of the enhanced TBP binding induced by destabilizing C-C mismatches when introduced at any of the 4 positions showing unstacking and destabilization in TBP-bound Watson-Crick DNA[15,16].**
(e) Destabilizing C-C mismatches enhance TBP binding when they are introduced at any of the four positions in the two base-pair steps (marked by red dotted rectangles) that are also destabilized in the TBP-bound Watson-Crick DNA[15,16]**.** The effect of the C-C mismatch at only one of these four positions (the last position in the 5'-TATAAAAG-3' site) was tested using the original SaMBA protocol, which is based on introducing single-base mutations in wild-type TF binding sites (in this case G-C to C-C). For the data shown here, we used a modified protocol to introduce double-base variations in TF binding sites by designing self-hybridizing DNA probes on the microarray (**Data File 1**).
(f) C-C mismatches do not enhance binding when introduced at any of the other positions in the TBP binding site.
(g) At all four positions in the two destabilized base-pair steps, mutations that rescue the Watson-Crick pairing do not have the same effect on TBP binding as the C-C mismatches.

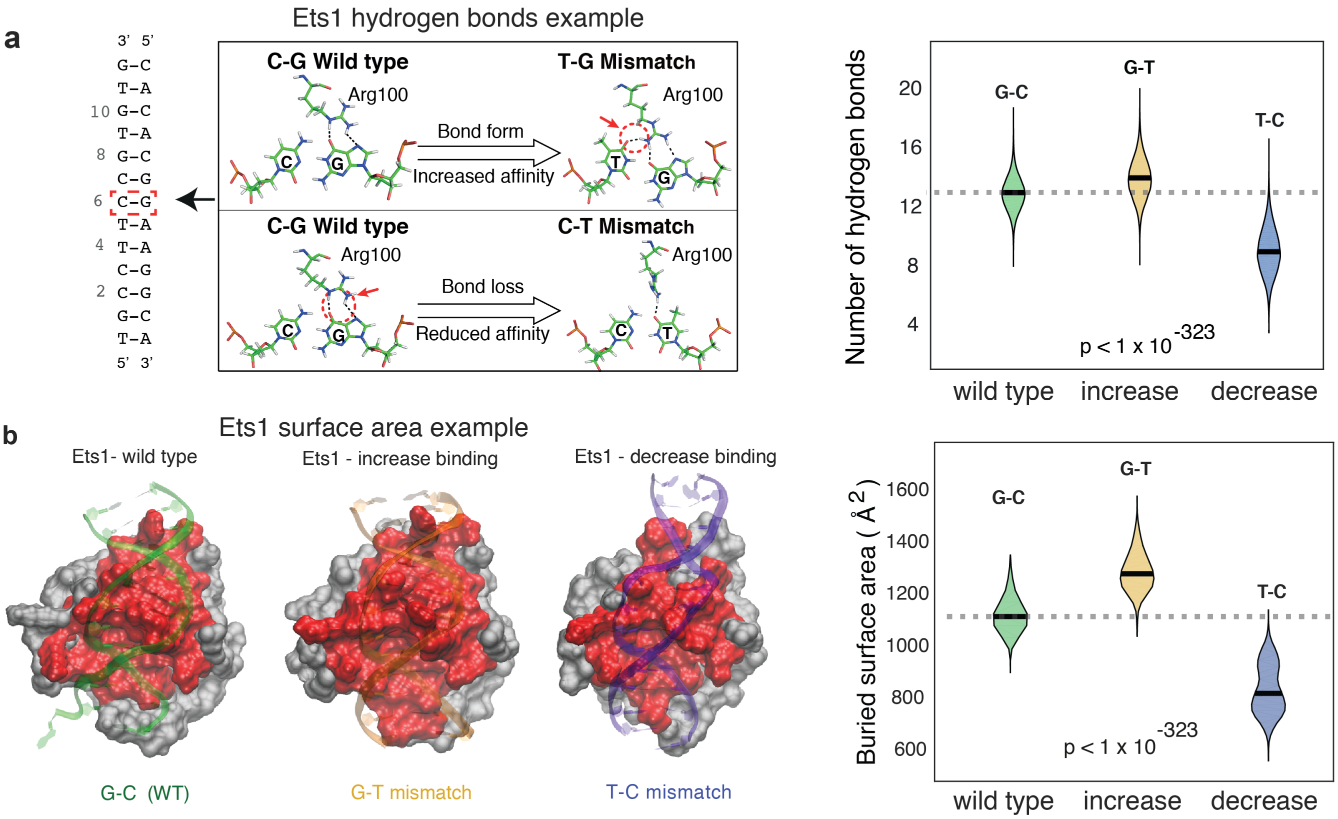


**Figure S10. Mismatch-induced changes in TF-DNA binding can also occur due to changes in direct readout.**
**(a) Mismatches in an Ets1 binding site result in changes in the number of protein-DNA hydrogen bonds** (based on MD simulations; Methods), which are correlated with changes in binding affinity. Left: representative snapshots showing the interaction between the C-G/T-G/C-T base pairs (red box in secondary structure) and Arg 100 (Arg 391 in 2NNY) on Ets1. Right: violin plots showing the distribution of the total number of H-bonds across the MD trajectory for different complexes (see Methods). Wilcoxon signed rank test p-value is shown.
**(b) Changes in hydrogen bonding in Ets1 are also accompanied by changes in the buried surface area** (see Methods), which are correlated with changes in binding. Left: TF is shown in gray and the buried surface area is shown in red. Right: violin plots show the buried surface area distribution in each trajectory. Wilcoxon signed rank test p-value is shown.
