## Supplementary material for "DNA mismatches reveal widespread conformational penalties in protein-DNA recognition": Methods and discussions

### Methods and Supplementary Discussion for DNA mismatches reveal widespread conformational penalties in protein-DNA recognition

Ariel Afek[1,2], Honglue Shi[3], Atul Rangadurai[4], Harshit Sahay[1], Hashim M. Al-Hashimi[3,4]*, Raluca Gordan[1,2,5]*

[1]Center for Genomic and Computational Biology, [2]Department of Biostatistics and Bioinformatics, [3]Department of Chemistry, [4]Department of Biochemistry, [5]Department of Computer Science, Department of Molecular Genetics and Microbiology, Duke University. * Corresponding authors.

#### Methods

##### Structural survey of standard Watson-Crick base pairs, distorted Watson-Crick base pairs in protein-DNA complexes, and mismatched base pairs in DNA

We performed a comprehensive survey of DNA base pair (bp) structures deposited in RCSB PDB[1]. X-ray crystal structures as well as NMR solution structures containing DNA were downloaded with their explicit PDB information including resolution, macromolecule type etc. from the RCSB website server on Aug 16$^{th}$, 2017. An in-house Python program was used to parse all structures using X3DNA-DSSR[2] into a searchable database (base-pair database) as described in previous studies[3]. The database contains all DNA base pairs in the biological unit with an accompanying list of structural parameters defining those base pairs. For all the following statistical analyses, crystal structures with resolution > 3 Å were excluded.

_Local base pair parameters:_ Base pair parameters—shear (Sx), stretch (Sy), stagger (Sz), buckle (κ), propeller twist (π), opening (σ), as well as C1′-C1′ distance of a given base pair—were computed using X3DNA-DSSR[2] as described previously[4]. The sign of shear (Sx) and buckle (κ) for all the A-T and G-C Watson-Crick base pairs were adjusted according to the index order of purine and pyrimidine (i.e. negative shear value indicates pyrimidine translates to major groove direction relative to purine). The base pair parameters of the bases with *syn* conformation (e.g. Hoogsteen base pairs) were not interpreted due to incorrect reference frame.

_Survey of standard Watson-Crick base pairs in B-DNA:_ To construct a dataset of standard Watson-Crick (WC) base pairs in B-DNA, the in-house Python program was used to identify all the canonical WC base pair structures in B-DNA environment following several criteria: (1) the canonical WC base pair is from only naked DNA structures without any protein or ligand bound, (2) base pairs with modified base are not considered, (3) the WC base pair is surrounded by at least two canonical WC base pairs from both sides in a DNA stem, and (4) non B-form DNA such as A-form or Z-form DNA were removed. A dataset containing a total of 903 A-T and 746 G-C standard WC bps was generated, and used to define the B-DNA envelope (**Table S1**).

_Survey of severely distorted Watson-Crick base pairs in protein-DNA complexes:_ To locate all distorted WC base pairs in the PDB database, the program was used to identify base pairs that satisfy the following criteria: (1) the distorted WC base pair is in a protein-DNA complex structure, (2) base pairs with modified bases are not considered, (3) the distorted WC base pair is surrounded by at least two canonical WC base pairs from both sides, and (4) the base pair has geometric parameters that are outliers relative to the B-DNA envelope: |shear (Sx)| > 1.5 Å or |stretch (Sy)| > 1.0 Å or |stagger (Sz)| > 1.5 Å or |buckle (κ)| > 30° or -35° < propeller (π) or propeller (π) > 15° or |opening (σ)| > 20° or |C1′-C1′ distance – (10.6 Å)| > 1.0 Å. The statistics of these distorted WC base pair are summarized in **Table S2.**

_Survey of DNA mismatches in DNA helical context:_ To survey the DNA mismatch structure and geometry, the program was also used to search and identify all possible single DNA mismatches (excluding modified bases) surrounded by at least two canonical WC base pairs from both sides, and were then subjected to manual inspection. Our survey finally identified a total of 44 G-T, 13 A-C, 12 G-A, 8 A-A, 9 G-G, 15 T-T, 6 C-T and 3 C-C mismatches within canonical DNA duplex context (**Tables S3,S4**). Please see Supplementary Discussion for a systematic analysis of DNA mismatch conformations.

#### Molecular dynamics (MD) simulations

All MD simulations were performed using AMBER ff99 force fields[5] with bsc0 corrections for DNA[6] and ff14SB corrections for proteins[7], and using standard periodic boundary conditions as implemented in the AMBER MD package[8]. The crystal waters in all structures were retained. The structures were then solvated using a truncated octahedral box of SPC/E[9] water molecules, with box size chosen such that the boundary was at least 10 Å away from any of the atoms of the free-DNA or protein-DNA complex using the leap module of the AMBER package. The default protonation states for protein residues that were assigned by leap were retained. $Na^+$ ions treated using the Joung Cheatham parameters[10] were then added to neutralize the charge of the system. The parameters used for the subsequent simulation setup are the same as those used in a prior study[4].

_MD simulations of free DNA with single mismatch:_ To simulate individual mismatches in free DNA, we started with two DNA duplexes: 5'- CTCTGCC**A**CGTGGGTCGT-3' and 5'-CACACG**G**AAGGCA-3'. We chose these sequences in part because they were also used SaMBA experiments for Myc/Max and Ets1; the bases of the DNA in the crystal structures (PDB IDs 1NKP and 2NNY) were mutagenized in silico to match those used in the binding experiments to obtain starting structures for the simulations. In the first sequence we focused on a parent A-T base pair (underlined), whereas in the second sequence we focused on a parent G-C base pair (underlined). We then performed single in-silico mutagenesis on either strand to create G-T, C-T, T-T, A-G, A-C, A-A mismatch for the parent A-T base pair in sequence 1, and C-C, G-G mismatch for the parent G-C base pair in sequence 2. For Hoogsteen mismatches like G(*syn*)-G, G-A(*syn*), one of the bases was manually rotated about the glycosidic bond by 180° to generate a *syn* conformation. Production runs of 500ns were carried out and extended to achieve convergence of the RMSD of the DNA if necessary. Based on insights obtained from the crystal structure survey (**Tables S3, S4**), the initial mismatch geometry was modeled as follows: (1) G-T and T-T mismatches were modeled as Wobble conformations; (2) C-T mismatch was modeled with two H-bonds (C:N3…T:H3-N3 and C:H4-H4…T:O4); (3) C-C mismatch was modeled with no H-bond but still stacking in the helix; (4) G-G mismatch was modeled as either G(*syn*)-G(*anti*) or G(*anti*)-G(*syn*) conformation; (5) G-A mismatch was modeled as either G(*anti*)-A(*anti*) or G(*anti*)-A(*syn*) conformation. No additional protonation of base was considered in these models.

_MD simulations of protein-DNA complexes:_ Starting structures corresponding to the Myc/Max and Ets1 systems were obtained from PDB entries 1NKP and 2NNY, respectively. For Ets1, the first ten residues of the unstructured N-terminal region were removed to get a starting structure for the simulations. Missing protein residues at the terminal ends for all systems were not modeled. For both systems, the bases of the DNA in the crystal structure were also mutagenized in silico to match those used in the binding experiments. Production runs of 200 ns (Myc/Max) or 500 ns (Ets1) were then carried out and extended to achieve convergence of the RMSD of the protein-DNA complex if necessary. For proteins bound to mismatched DNA sites, simulations were performed only for C-T and G-T mismatches, as these mismatches exist in one stable conformation that can be reliably modeled using MD force fields. Other mismatches either have multiple conformations, and MD simulations cannot capture exchanges between these conformations, or involve protonation/de-protonation, which is also not modeled well by MD.

#### Protein expression and purification

_Ets1, Elk1, Gabpa, Runx1, Six6, Ap2a, Gata1:_ Gateway-compatible clones containing the full-length genes for human Ets1, Elk1, Gabpa, Runx, Six6, Ap2a, and Gata1 were purchased from GeneCopoeia and the genes were transferred into pDEST15 using the LR Clonase reaction (Life Technologies). This vector enables the production of N-terminally GST-tagged proteins. Cells were grown in LB broth to an OD600 of 0.4 to 0.6 and then expression was induced with IPTG at 30 to 37 °C. Pelleted cells were frozen, stored at -20 °C, and thawed cells were lysed with lysozyme. GST-tagged protein was purified from the soluble portion of the lysate using GST resin (GE Healthcare) according to manufacturer's instructions.

_Cbf1:_ *S. cerevisiae* Cbf1 was overexpressed in *E. coli* BL21 (DE3) cells (New England BiolLabs) and purified by as previously described[11].

_E2f1:_ The Life Technologies Gateway cloning system[2] was used to insert full-length human E2f1 genes into the pET-60 destination vector with a C-terminal GST tag. Cells were grown in LB culture to an OD600 of 0.4, then protein expression was induced with 1mM IPTG at 30 °C for 2hrs. Pelleted cells were frozen, then thawed cells were lysed in

PBS lysis buffer for two hours at 4 °C while gently rocking. The lysate was centrifuged, and the protein was recovered from the soluble lysate using a GE GSTrap FF GST tag affinity column according to manufacturer's instructions.

_Myc/Max/Mad:_ Full-length human Myc (Myc), Max, and Mad (Mad1) proteins with C-terminal 6xHis tags, as well full-length untagged Max protein were generously provided by Peter Rahl and Richard Young (Whitehead Institute and MIT). The proteins were expressed in bacteria and purified as described in a prior study[3]. As Myc requires heterodimerization with Max to bind DNA efficiently, all Myc SaMBA experiments were performed using both Myc and Max on the same microarray, using a 5 times higher concentration of Myc compared with Max to ensure that mostly Myc:Max heterodimers, and not Max:Max homodimers, are formed. Similarly, all Mad SaMBA experiments were performed using both Mad and Max on the same microarray, with a 5 times higher concentration of Mad.

_Egr1:_ The Egr1 zinc-finger protein (human Egr1 residues 335− 423) was generously provided by Junji Iwahara (University of Texas Medical Branch). The protein was expressed in *E. coli* strain BL21 (DE3) and purified as previously described[4]. The binding buffer for this protein was: 10 mM Tris-HCl (pH 7.5), 150 mM KCl, and 0.2 μM $ZnCl_2$, 2% (wt/vol) milk ,51.3 ng/μl salmon testes DNA, 0.2 μg/μl bovine serum albumin.

_p53:_ Full-length recombinant human p53 protein with N-terminal GST tag was purchased from Abcam (ab43615).

_TBP:_ Full-length recombinant human TBP protein with HIS tag was purchased from Excellgen (catalog#: RP-54). The binding buffer for this protein was 10 mM HEPES, 70 mM KCl, 10 mM $MgCl_2$, 1 mM EDTA, 2% (wt/vol) milk, 51.3 ng/μl salmon testes DNA, 0.2 μg/μl bovine serum albumin.

_CTCF:_ Full-length recombinant human CTCF protein with N-terminal GST tag was purchased from Abnova (catalog #:H00010664). The binding buffer for this protein was PBS with 5 mM $MgCl_2$, 0.1 mM $ZnSO_4$ [5] , 2% (wt/vol) milk , 51.3 ng/μl salmon testes DNA, 0.2 μg/μl bovine serum albumin.

_Creb1/Crem/Atf1:_ Full-length recombinant human Creb1/Crem/Atf1 proteins with N-terminal HIS tags were purchased from Origene (catalog #: TP760318/ TP760397/TP721193). Binding buffer for Creb1/Crem was 25 mM Tris-HCl (pH 7.4), 1 mM DTT, 0.5 mM EDTA, 2% glycerol, 5 mM $MgCl_2$, 50 mM KCl, 25 mM Boric Acid. Binding buffer for Atf1 was 50 mM Potassium Acetate, 20 mM Tris-Acetate, (pH 7.9) 10 mM Magnesium Acetate, 1 mM DTT. Both buffer solutions also contained 2% (wt/vol) milk, 51.3 ng/μl salmon testes DNA, 0.2 μg/μl bovine serum albumin.

_Stat3 and RelA:_ Mouse phosphorylated Stat3 (residues 127-716) with 6xHis tag, and Human RelA (residues 20-290) were generously provided by Dr. Eyal Arbely (Ben-Gurion University of the Negev). Binding buffer for RelA was: 6mM HEPES, 80 mM KCl, 0.5 mM EDTA, 2% (wt/vol) milk , 51.3 ng/μl salmon testes DNA , 0.2 μg/μl bovine serum albumin. RelA rabbit polyclonal antibody was purchased from Origene (Catalog #: TA890002).

#### SaMBA library design and measurements

Saturation Mismatch Binding Assays (SaMBA) were performed as follows. First, four custom DNA libraries (v1-v4), each containing ~15,000 single-stranded 60-base oligonucleotides, were designed computationally based on TF binding site sequences for 21 TF proteins (**Table S5; Data File 1**). The binding site(s) for each TF were selected based on published data showing specific TF binding to these sites. Sites were selected to contain central 8-mers with protein-binding microarray (PBM) enrichment scores (E-score) of 0.35 or higher, which is indicative of specific protein binding[12,13]. Given that universal PBM data was not available for CTCF, p53 and RelA, for these proteins we selected the strongest binding sites based on the DNA binding motif reported in the Jaspar databse[14] (for CTCF), the binding data reported in[15] (for p53), and the binding data reported in[16] (for RelA).

Each DNA library was designed to contain multiple replicates (8-20) of both wild-type (WT) binding site sequences and all possible single base variants of the same sites. For SaMBA library "v1", we used 14 replicate spots for the wild-type sequence and 8 replicate spots for each mismatch. For SaMBA libraries "v2", "v3", and "v4" we used 20 replicate spots for the wild-type sequence and 10 replicate spots for each mismatch. The DNA libraries were commercially synthesized on DNA microarray chips (Agilent). Next, double stranded DNA binding sites were generated on the chip by hybridization with the WT reverse complement oligonucleotides in solution (variant complements were absent from the hybridization solution). The complement oligonucleotide solution was composed of ~2.5 μM (large excess) HPLC-purified oligonucleotides, and ~0.25 μM HPLC-purified oligonucleotides of identical sequence with FAM/Cy3 fluorophores attached (Integrated DNA Technologies). The absence of perfect complements in the solution for the variant sequences on the chip ensured successful hybridization with the WT complements for both the variants and WT sequences present on the chip. The small fraction of fluorescently labeled oligonucleotides allowed us to assess the successful formation of the mismatched duplexes on the chip (**Fig. S4**).

The reaction buffer mixture for the hybridization step was 100 μl 10x reaction buffer (260 mM Tris-HCl, pH 9.5, 65 mM $MgCl_2$) in a total volume of 1000 μl, similar to a prior study[12]. The chip (Agilent) was incubated with reaction mixture in a hybridization oven using a pre-warmed stainless-steel hybridization chamber and gasket cover slip. After a 5-hours incubation (85 °C for 10 min, 75 °C for 10 min, 65 °C for 60 min, 60 °C for 120 min, and 55 °C for 100 min), the hybridization chamber was disassembled in a glass staining dish in 500 ml phosphate buffered saline (PBS) / 0.01% Triton X-100 at 37°C. The chip was transferred to a second staining dish, washed for 10 min in PBS / 0.01% Triton X-100 at 37°C, washed once more for 3 min in PBS at room temperature, similar to a prior study[12]. The fluorescent signal (Cy3/FAM) of hybridized oligonucleotides was scanned and measured using a GenePix 4400A microarray scanner to confirm that the hybridization was successful and reproducible, and that no significant cross-hybridization occurs (**Fig. S4**, **Fig. 1e; Data File 1**).

Protein binding reactions were performed in the same conditions as used in PBM protocols[12]. The binding buffer, unless otherwise specified, was a 185-µl solution containing PBS / 2% (wt/vol) milk / 51.3 ng/µl salmon testes DNA (Sigma) / 0.2 µg/µl bovine serum albumin (New England Biolabs). Pre-incubated protein binding mixtures were applied to individual chambers and incubated for 1 h with the double-stranded DNA chip. The chips were washed once with PBS / 0.5% (vol/vol) Tween-20 for 3 min, and then once with PBS / 0.01% Triton X-100 for 2 min. After the protein incubation and washing steps, Alexa647-conjugated GST antibody (Invitrogen),Penta·His Alexa647-conjugated antibody (Qiagen), or Goat anti-Rabbit IgG Alexa647 antibody (ThermoFisher) in PBS / 2% milk were applied on the chip for 1 h at room temperature. Washing steps after each incubation step were performed in Coplin jars at room temperature, on a shaker at 125 r.p.m. The fluorescent signal (Alexa647) of bound TFs for each DNA spot was measured using a GenePix 4400S microarray scanner. Multiple replicates of each sequence were used to quantitatively compare the binding signals between sequences, and to statistically assess the significance of binding differences using a two-sided Wilcoxon-Mann-Whitney test (**Fig. S5**).

#### Fluorescence anisotropy experiments for p53

The p53 florescence anisotropy measurements were performed at room temperature with BMG Clariostar multimode plate reader, and Corning low volume 384-well black flat bottom polystyrene NBS microplate. 5' 6-FAM labeled, HPLC-purified oligonucleotides of both 'mismatched' and 'WT' duplexes were obtained from Integrated DNA Technologies (IDT). The buffer conditions for the binding experiments were PBS, 10 % v/v glycerol 5 mM DTT, and 0.1 mg/ml BSA. Florescence anisotropy signal was measured for 1nM 6-FAM p53 DNA binding site ('WT' and 'mismatched'), in 17 different p53 concentrations, with 3 replicates for each concentration. The fluorescence anisotropy signal was fit to the two-site equilibrium binding model: $K_d = \frac{[P]^2[DNA]}{[P_{2\circ}DNA]}$ , where [P] is p53 dimer concentration, and [DNA] is the DNA concentration. Since [DNA]<<[P], using $[P]_0$=[P] approximation in the above equations gives the following expression for the observed anisotropy: $r_{obs} = r_0 + \frac{(r_{bound} - r_0) \times [P]_0{}^2}{Kd + [P]_0{}^2}$ where $r_{obs}$ is anisotropy observed for each p53 concentration, $r_0$ is the initial anisotropy value of the unbound DNA and $r_{bound}$ is the anisotropy upon binding of p53 to DNA. The $K_d$ shown is the overall dissociation constant for the entire binding process.

#### Protein binding microarray (PBM) experiments

For measuring the effect of mutations on transcription factor binding, we used the standard PBM protocol[12], but with a custom DNA library containing TF binding sites of interest and variants of these sites (**Data File 1**).

#### Nuclear magnetic resonance (NMR) experiments

Single strands corresponding to the WT p53 binding site and single mismatches sites were purchased from IDT with standard desalting purification. After re-suspension in water, the equimolar amounts of single strands were mixed together to form the WT and T-T mismatch-containing duplexes. Concentrations were measured using a Nanodrop 3000, with the extinction coefficients for single and double strands obtained using the ADT bio oligo calculator. The duplexes were annealed by heating to 95 °C for 5 min and cooling at room temperature for ~1 hr. They were then exchanged into NMR buffer (15 mM Sodium phosphate, 25 mM Sodium Chloride, 0.1 mM EDTA, pH 6.9) using centrifugal concentrators. 500 μL solutions containing 10% $D_2O$ were prepared for collecting proton 1D spectra. An analogous protocol was used for preparing A6DNA duplexes containing A-T, m1A-T, T-T and C-T base pairs with the m1A containing single strand being purchased from Yale Keck Oligonucleotide Synthesis Facility with HPLC purification. Duplex samples containing 10% $D_2O$ after buffer exchange were lyophilized into 100% $D_2O$. Assignments of the sugar resonances were performed using a combination of 2D $^1$H-$^1$H NOESY, 2D $^1$H-$^1$H TOCSY and 2D $^1$H-$^{13}$C HSQC experiments.

#### Structural analyses of mismatches that enhance TF binding

Structures of protein-DNA complexes are available in PDB for 12 of the 21 TFs examined by SaMBA (**Table S6**). For one additional protein, Stat3, we did not use the PDB structure in our analyses due to inconsistencies in the PDB data file. When multiple structures were available for the same TF, we chose the one with the DNA sequence most similar to the one tested in SaMBA (**Table S6**). For the selected structures, we then focused on the regions that are identical between the crystal/NMR structure and the SaMBA sequence (underlined in **Table S6**), but ignoring the positions that are at the terminal ends in the crystal structures. We identified 21 positions at which we found increased TF binding, due to a total of 29 mismatches (for some positions we found several mismatches that lead to increased protein binding levels). For these 21 positions, we examined the protein-bound DNA structure and the mismatch structure to determine whether the mismatch is inducing structural features that mimic the bound geometry. We found mimicry in the case of 15 of the 29 mismatches (i.e. 52%) (**Table S7**). Since mutating a base in contact with the protein can lead to a loss or gain of protein-DNA contacts, we also re-examined the structures after removing from consideration all mismatches at bases in contact with the protein (according to the protein-DNA structure; see **Table S7**). We found 11 cases of mimicry out of 22 total mismatches (i.e. 50%). To further separate the structural effect from potential sequence effect, we also performed an analysis where we removed from consideration all mismatches for which the rescue Watson-Crick mutation also increases TF binding affinity (based on mutation binding data, **Fig. S8**, and the DNA binding motifs of the TF proteins, **Fig. S6**). In this analysis, we found 14 cases of mimicry out of 28 total mismatches (i.e. 50%). Thus, about half of the mismatches that enhance TF binding induce deformations similar to those observed in the bound DNA structure.

#### Inter-helical Euler angle calculation

We used a well-established inter-helical Euler angle approach to quantify DNA local kinking angles with the bending magnitude ($\beta_h$, $0^o \leq \beta_h \leq 180^o$), the bending direction ($\gamma_h$, $-180^o \leq \gamma_h \leq 180^o$), and the helical twist ($\zeta_h$, $-180^o \leq \zeta_h \leq 180^o$) of two helices (H1 and H2) across a given base pair junction[3,4,17,18]. The junction can be a Watson-Crick base pair, Hoogsteen base pair, or any mismatched pair. In this approach, two idealized B-form DNA helices constructed by 3DNA[19] and containing 3 base pairs are superimposed to H1 and H2, respectively, yielding an relative orientation of H1 to H2 with parameters ($\beta_h$, $\gamma_h$, $\zeta_h$). All calculations with poor alignment to the idealized helix (RMSD > 2 Å for only sugar and backbone atoms)[17] were omitted from analysis. To visualize the bending magnitude and direction of H2 relative to H1, we plot the average $\beta_h$ and $\gamma_h$ for a given base pair junction in a polar coordinate in which the radial axis represents the $\beta_h$ ($0^o \leq \beta_h \leq 60^o$) and the angular axis represents the $\gamma_h$ ($-180^o \leq \gamma_h \leq 180^o$).

To evaluate the similarity of inter-helical Euler angle distribution between free and bound DNA in the MD simulation (**Fig. 3f**), we used a well-established one-dimensional REsemble analysis[20] to quantify the overlap between free and bound DNA kinking angle distribution over K increment bin sizes from 5° to 360°. The output Ω value (from 0 to 1) represents the similarity between free and bound distributions in which 0 indicates perfectly identical and 1 indicates no similarity.

Using the approach described above, we analyzed and compared the changes in kinking angle induced by the G-T versus C-T mismatches in a Myc binding site (**Fig. 3f)**. We chose these mismatches because: (i) they have opposite effects on Myc binding levels (with G-T increasing Myc binding by ~2-fold and C-T decreasing Myc binding by ~5-

fold), and (ii) these mismatches have well-defined base-pair geometries at neutral pH, compared to all other mismatches (**Fig. S2a**).

##### H-bond and buried surface area analyses

The number of hydrogen bonds between the protein and DNA (**Fig. S10a**) were calculated using the hbond program of the CPPTRAJ[21] utility of AMBER. A heavy atom donor (Nitrogen/Oxygen/Sulphur) with an attached hydrogen was defined as forming a hydrogen bond with an acceptor heavy atom (Nitrogen/Oxygen/Sulphur) if the donor-acceptor distance was less than 3 Å and the acceptor-hydrogen-donor angle was greater than 135°. The calculation was performed by considering the protein and DNA atoms as donors/acceptors separately and adding the resulting number of hydrogen bonds. The number of hydrogen bonds was then averaged over all conformers in the MD trajectory for a given system.

The buried surface area for a given conformation of a protein-DNA complex (**Fig. S10b**) was defined as the difference between the sum of surface areas of the protein and DNA when considered in isolation, and the protein-DNA complex. The computation of the surface area was performed using the molsurf program in the CPPTRAJ [21] utility of AMBER, with a probe radius of 1.4 Å. The surface area was averaged over all conformers from the MD simulations to obtain the buried surface area for the system. All the residues of the protein and DNA were considered for the calculation; similar results were obtained for the Ets1 system on the exclusion of the terminal ends of the DNA (data not shown).

#### <u>Supplementary Discussion</u>

##### Systematic analysis of DNA mismatch conformations

Mismatches are proposed to adopt multiple conformations that are undergoing exchange in solution (**Figure S2a**). For example, G(*syn*)-G(*anti*) Hoogsteen mismatches exist in dynamic equilibrium with G(*anti*)-G(*syn*); G-T wobble mismatches exist in dynamic equilibrium with Watson-Crick-like bps stabilized by rare tautomeric and anionic bases; G-A mismatches adopt multiple conformations involving protonated adenine and either *syn* or *anti* bases.

To systematically analyze the ensemble behavior of all mismatches and examine their conformational penalty on protein-DNA recognition, we performed MD simulations on unbound DNA with different single mismatches (Methods). Base pair parameters and the C1′-C1′ distance are used to parameterize the conformation of mismatches. Note that: (1) transient and rare species, such as the tautomeric form of the G-T mismatch, are ignored in this analysis; (2) due to the inability of glycosidic bond rotations to occur under timescales conventionally accessible by MD simulations , different Hoogsteen-type mismatches, i.e. G(*syn*)-G(*anti*) and G(*anti*)-G(*syn*), are modeled and simulated separately; (3) protonated or anionic base pairs, such as $A^+$-C, $C^+$-C and G(syn)-$A^+$(anti), are ignored in this analysis for simplicity since they are highly dependent on pKa and pH (Methods). A summary descriptions of the ensemble behavior of different mismatches in the unbound DNA MD simulation are presented below and in **Fig. S2b**.

<u>*Purine-pyrimidine mismatches*</u>: Purine-pyrimidine mismatches consist of G-T and A-C. The G-T mismatch remains in a stable wobble geometry with shear around -2 Å, accompanied by a slight constriction (stretch and C1′-C1′ distance) during the MD simulation, compared to A-T and G-C base pairs. The A-C mismatch remains mostly as one H-bonding geometry with C translated to the major groove (similar to G-T), whereas in a prior computational investigation, A-C mismatches exist dynamic equilibrium of either A or C translating to major groove[22]. Interestingly, in our MD simulation, A-C mismatch is slight positively buckled.

<u>*Pyrimidine-pyrimidine mismatches*</u>: Pyrimidine-pyrimidine mismatches consist of T-T, C-T, and C-C. T-T mismatch remains in a wobble geometry with shear around ±2 Å. In contrast to the G-T wobble mismatch, the T-T mismatch exists in rapid dynamic equilibrium of both inter-converting wobble geometry with either one of the T translating to the minor groove direction, as shown in the MD simulation. Despite this rapid dynamic equilibrium, the T-T mismatch is still constricted with C1′-C1′ distance of 8-9.5 Å during the MD simulation. Similar to T-T, the C-T mismatch is also constricted with two H-bonds stably formed for most of the time. However, C-T can transiently adopt a high-energy conformation with only one H-bond formed, and it is partially open (C1′-C1′ distance around 10 Å), potentially due to the repulsion between T-O2 and C-O2. The entire C-T MD trajectory is comprised of approximately 5% of these high-energy species. The C-C mismatch is partially constricted with C1′-C1′ distance around 9.8 Å, due to its unstable one

H-bonding geometry. All the pyrimidine-pyrimidine mismatches were still stacked in the helix without swinging outside the helix during the MD simulation.

Purine-purine mismatches: Purine-purine mismatches consist of G-G, G-A, and A-A. The G-G mismatch basically only exists as a mixture of inter-converting G(*syn*)-G(*anti*) conformations in a sequence-dependent manner[23]. During the MD simulation, G(*syn*)-G(*anti*) does not experience syn-anti inter-converting exchange. The C1′-C1′ distance of G(*syn*)-G(*anti)* is around 11.2-11.5 Å, which is larger than for the canonical G-C base pair. G(*anti*)-A(*anti*) and G(*syn*)-A(*anti*) geometries were considered for the G-A mismatch. G(*anti*)-A(*syn*) is partially expanded, with C1′-C1′ distance around 11.5 Å, whereas G(*anti*)-A(*anti*) is strongly stretched, with a large C1′-C1′ distance around 12.8 Å. The A-A mismatch adopts an unstable stretched A(*anti*)-A(*anti*) geometry, with only one H-bond.

We also performed an independent analysis of DNA mismatch structures within helical context, from a PDB structural survey (Methods). A detailed summary of the structural survey is listed in **Table S3**. We note that: (1) we only consider entries with a single mismatch embedded by two Watson-Crick base pairs; and (2) modified bases were excluded from the survey (Methods). Importantly, most of T-T and C-T mismatches are constricted (C1′-C1′ distance < 9.5 Å), with only one exception (PDB: 2LL9), whereas all G-T and A-C mismatches are not constricted (C1′-C1′ distance > 10.0 Å). The C-C mismatch has only three entries, with only one of them constricted (PDB: 2NL8**.** Purine-purine mismatches (G-A, A-A, G-G) are also not constricted, consistent with a previous survey about purine-purine mismatches[23]. Additionally, consistent with MD simulation, G-T, A-C and T-T mismatches adopt mostly wobble conformations.

#### Potential roles for TF binding to mismatches in the cell

TF binding to DNA is a key determinant of gene regulation. Alterations in TF binding sites can affect cell functions and lead to human disease[24]. A well-characterized example is that of TFs from the ETS family, which selectively bind to the *TERT* promoter and activate this gene in cancer cells, due to single base-pair mutations that create high affinity ETS sites[25,26]. Remarkably, using SaMBA we observed several mismatches that also forms high affinity ETS sites (**Fig. S5, S6**). Beyond ETS proteins, human cells express hundreds of TF proteins, which collectively bind to hundreds of thousands of genomic sites[27]. Mismatch in these sites have the potential to alter TF binding and affect cellular activity.

The cellular effects of TF binding to mismatched DNA will depend on both the TF proteins (their abundance and affinities for mismatches), and the mismatches (their abundance and correction efficiency, which varies widely across the genome[28]). Several recent studies suggested that TFs could rapidly bind nascent DNA strands[29]; nevertheless, the fast repair of most replication errors significantly reduces the chance of TFs recognizing mismatches that arise due to errors during replication. On the other hand, the repair of mismatches that are formed during genetic recombination, or during gap-filling DNA synthesis, is slower and less efficient, with T-T and C-C mismatches repaired with very low efficiency[28,30]. Furthermore, spontaneous deamination is common and estimated to occur 100-500 times per cell per day in humans[31]. G-T mismatches that are generated by the deamination of 5-meC are not repaired by the mismatch repair pathway and have considerably lower repair efficiency[31]. In such cases, given that many TFs are bound all across the genome to regulate a variety of cellular processes, it is reasonable to expect that TFs would be functionally affected by the presence of mismatches. In tumor cells with defects in DNA mismatch repair, the effect of mismatches on TFs is expected to be even larger than in healthy cells.

TFs bound to DNA mismatches in the genome could also affect damage correction and lead to the formation of genetic mutations. Several recent studies hypothesized that TFs are likely playing a role in shaping the mutational landscape of eukaryotic genomes by interfering with DNA repair and replication[29,32,33]. In particular, mismatches bound with high affinity by TFs may be harder to recognize by repair enzymes[33] and harder to correct during strand replacement by Pol-$\delta$[29] (**Fig. S3d**). Our study provides a platform to map which specific mismatches attract TFs and could become roadblocks for DNA repair and replication, thus contributing to mutagenesis and even becoming potential mutational "hotspots" in the genome.

Finally, TFs binding to mismatched DNA could also play useful roles in damage recognition and the cellular response to damage. For example, p53 is known to sense and respond to DNA lesions by different mechanisms[34], and to regulate cellular processes such as cell cycle arrest, apoptosis and DNA repair. Since other TFs are also involved in these processes, understanding the interplay between TFs and mismatched DNA could provide insights into DNA damage response mechanisms in the cell.
