## Supplemental tables for "DNA mismatches reveal widespread conformational penalties in protein-DNA recognition"

### Supplementary Tables for
#### DNA mismatches reveal widespread conformational penalties in protein-DNA recognition

Ariel Afek[1,2], Honglue Shi[3], Atul Rangadurai[4], Harshit Sahay[1], Hashim M. Al-Hashimi[3,4]*, Raluca Gordan[1,2,5]*

[1]Center for Genomic and Computational Biology, [2]Department of Biostatistics and Bioinformatics, [3]Department of Chemistry, [4]Department of Biochemistry, [5]Department of Computer Science, Department of Molecular Genetics and Microbiology, Duke University. * Corresponding authors.

**Table S1. Defining the B-DNA envelope: summary statistics of base pair parameters (mean, maximum value, minimum value, and standard deviation) for base pairs in B-DNA, obtained from a comprehensive survey of structures deposited in RCSB PDB (Methods).**

**(a) A-T/G-C for both X-ray and NMR**

| parameter | mean | maximum | minimum | standard deviation |
|---|---|---|---|---|
| shear | -0.08 | 1.64 | -2.15 | 0.36 |
| stretch | -0.15 | 0.73 | -0.82 | 0.15 |
| stagger | 0.09 | 1.66 | -1.43 | 0.29 |
| buckle | 1.88 | 27.96 | -29.69 | 7.53 |
| propeller | -10.13 | 31.66 | -37.01 | 8.18 |
| opening | -0.44 | 23.94 | -18.07 | 3.84 |
| C1′- C1′ distance | 10.6 | 12.5 | 9.2 | 0.23 |

**(b) A-T/G-C for X-ray**

| parameter | mean | maximum | minimum | standard deviation |
|---|---|---|---|---|
| shear | -0.03 | 1.64 | -2.15 | 0.37 |
| stretch | -0.14 | 0.73 | -0.82 | 0.17 |
| stagger | 0.1 | 1.12 | -1.43 | 0.25 |
| buckle | 2.33 | 22.55 | -29.69 | 7.07 |
| propeller | -11.92 | 14.49 | -36.65 | 6.55 |
| opening | 0.44 | 23.94 | -18.07 | 4 |
| C1′- C1′ distance | 10.56 | 12.5 | 9.7 | 0.24 |

**(c) A-T/G-C for NMR**

| parameter | mean | maximum | minimum | standard deviation |
|---|---|---|---|---|
| shear | -0.12 | 1.23 | -1.29 | 0.34 |
| stretch | -0.17 | 0.26 | -0.56 | 0.12 |
| stagger | 0.08 | 1.66 | -1.33 | 0.33 |
| buckle | 1.38 | 27.96 | -29.61 | 7.98 |
| propeller | -8.11 | 31.66 | -37.01 | 9.29 |
| opening | -1.43 | 14.9 | -15.14 | 3.39 |
| C1′- C1′ distance | 10.65 | 11.3 | 9.2 | 0.21 |

#### Table S2. Summary of extremely distorted Watson-Crick base pairs in protein-DNA complexes

| Cutoff [a] | Method | Number of occurrences | Number of PDB entries |
|---|---|---|---|
| Shear (Sx) < -1.5 Å OR > 1.5 Å | X-ray | 490 | 196 |
| | NMR | 4 | 3 |
| Stretch (Sy) < -1.0 Å OR > 1.0 Å | X-ray | 84 | 63 |
| | NMR | 4 | 3 |
| Stagger (Sz) < -1.5 Å OR > 1.5 Å | X-ray | 213 | 83 |
| | NMR | 14 | 11 |
| Buckle (κ) > 30° OR < -30° | X-ray | 260 | 126 |
| | NMR | 8 | 6 |
| Propeller (π) < -35° OR > 15° | X-ray | 322 | 139 |
| | NMR | 36 | 20 |
| Opening (σ) < -20° OR > 20° | X-ray | 224 | 114 |
| | NMR | 10 | 8 |
| C1′-C1′ distance < 9.6 Å OR > 11.6 Å | X-ray | 183 | 104 |
| | NMR | 8 | 7 |

a. Please see Methods for details.

#### Table S3. Structural survey of DNA mismatches

| Mismatch | PDB | Method | Macromolecule Type [a] | Resolution (Å) | Base pair ID [b] | Notes | Shear (Å) | C1′-C1′ distance (Å) |
|---|---|---|---|---|---|---|---|---|
| AA | 1DMU | X-ray | Protein#DNA | 2.2 | 1dmu_AA_F_DA_0_F_DA_0 | Anti-Anti | 0 | 11.107 |
| AA | 1OH6 | X-ray | Protein#DNA | 2.4 | 1oh6_AA_E_DA_9_F_DA_22 | Distorted Syn-Anti | 2.031 | 13.239 |
| AA | 1SVC | X-ray | Protein#DNA | 2.6 | 1svc_AA_D_DA_10_D_DA_10 | Anti-Anti | 0 | 10.867 |
| AA | 2A3V | X-ray | Protein#DNA | 2.8 | 2a3v_AA_E_DA_33_F_DA_8 | Anti-Anti | 2.043 | 12.968 |
| AA | 4IZZ | X-ray | Protein#DNA | 2.5 | 4izz_AA_C_DA_0_D_DA_0 | Anti-Anti | 0.639 | 11.355 |
| AA | 4J00 | X-ray | Protein#DNA | 3 | 4j00_AA_C_DA_13_D_DA_13 | Anti-Anti | -0.933 | 11.806 |
| AA | 4RB1 | X-ray | Protein#DNA | 2.75 | 4rb1_AA_D_DA_615_D_DA_615 | Anti-Anti | 0 | 10.524 |
| AA | 5BMZ | X-ray | Protein#DNA | 3 | 5bmz_AA_E_DA_0_F_DA_0 | Anti-Anti | 0.083 | 10.789 |
| AC | 1D99 | X-ray | DNA | 2.5 | 1d99_AC_A_DA_4_B_DC_21 | Wobble | -1.713 | 10.438 |
| AC | 1D99 | X-ray | DNA | 2.5 | 1d99_AC_A_DC_9_B_DA_16 | Wobble | 2.166 | 10.192 |
| AC | 1OH5 | X-ray | Protein#DNA | 2.9 | 1oh5_AC_E_DC_9_F_DA_22 | Distorted syn-anti | 0.225 | 11.863 |
| AC | 3POV | X-ray | Protein#DNA | 2.5 | 3pov_AC_C_DC_12_D_DA_29 | Distorted syn-anti | -1.199 | 10.882 |
| AC | 3TAP | X-ray | Protein#DNA | 1.65 | 3tap_AC_B_DC_27_C_DA_6 | Tautomer | 0.768 | 10.305 |
| AC | 3TAQ | X-ray | Protein#DNA | 1.65 | 3taq_AC_B_DC_26_C_DA_7 | Wobble | 2.128 | 10.551 |
| AC | 3TAR | X-ray | Protein#DNA | 1.6 | 3tar_AC_B_DC_29_C_DA_9 | Wobble | 2.056 | 10.422 |
| AC | 4RDM | X-ray | Protein#DNA | 2.7 | 4rdm_AC_C_DA_6_D_DC_28 | Tautomer | -0.548 | 10.494 |
| AC | 5D9I | X-ray | Protein#DNA | 1.7 | 5d9i_AC_X_DC_8_Y_DA_15 | Single h-bond | -1.376 | 11.298 |
| AC | 5FW3 | X-ray | Protein#DNA#RNA | 2.7 | 5fw3_AC_C_DC_4_D_DA_9 | Tautomer? | 0.572 | 10.531 |
| AC | 1N1N | NMR | DNA | null | 1n1n_AC_A_DC_5_B_DA_16 | Wobble? | 1.535 | 10.23 |
| AC | 2MO7 | NMR | DNA | null | 2mo7_AC_A_DA_4_B_DC_21 | Wobble? | -2.266 | 10.75 |
| AC | 2MO7 | NMR | DNA | null | 2mo7_AC_A_DC_9_B_DA_16 | Wobble? | 2.279 | 10.759 |
| AG | 111D | X-ray | DNA | 2.25 | 111d_AG_A_DA_4_B_DG_21 | G(syn).A(anti) | 0.235 | 11.006 |
| AG | 111D | X-ray | DNA | 2.25 | 111d_AG_A_DG_9_B_DA_16 | G(syn).A(anti) | -0.546 | 10.607 |
| AG | 112D | X-ray | DNA | 2.5 | 112d_AG_A_DG_4_B_DA_21 | G(anti).A(syn) | -0.124 | 10.809 |
| AG | 112D | X-ray | DNA | 2.5 | 112d_AG_A_DA_9_B_DG_16 | G(anti).A(syn) | -0.58 | 10.574 |
| AG | 1DNM | X-ray | DNA | 2.5 | 1dnm_AG_A_DA_4_B_DG_21 | G(anti).A(syn) | -0.187 | 11.386 |
| AG | 1DNM | X-ray | DNA | 2.5 | 1dnm_AG_A_DG_9_B_DA_16 | G(anti).A(syn) | 0.306 | 11.459 |
| AG | 4R22 | X-ray | Protein#DNA | 2.6 | 4r22_AG_G_DA_5_G_DG_15 | G(anti).A(anti) | 0.42 | 11.872 |
| AG | 4R22 | X-ray | Protein#DNA | 2.6 | 4r22_AG_G_DG_15_G_DA_5 | G(anti).A(anti) | -0.42 | 11.872 |
| AG | 4R24 | X-ray | Protein#DNA | 2.25 | 4r24_AG_G_DA_5_G_DG_15 | G(anti).A(anti) | 0.293 | 11.647 |
| AG | 4R24 | X-ray | Protein#DNA | 2.25 | 4r24_AG_G_DG_15_G_DA_5 | G(anti).A(anti) | -0.293 | 11.647 |
| AG | 5DBB | X-ray | Protein#DNA | 2.25 | 5dbb_AG_T_DA_13_P_DG_4 | G(anti).A(syn) | 0.314 | 10.79 |

| | | | | | | | | |
|---|---|---|---|---|---|---|---|---|
| AG | 1ONM | NMR | DNA | null | 1onm_AG_A_DA_6_B_DG_17 | G(anti).A(anti) | -0.323 | 12.529 |
| CC | 2NL8 | X-ray | Protein#DNA | 2.3 | 2nl8_CC_W_DC_19_W_DC_19 | water mediated | 0 | 10.76 |
| CC | 3ZVN | X-ray | Protein#DNA | 2.15 | 3zvn_CC_F_DC_3_H_DC_3 | Wobble | 1.62 | 8.594 |
| CC | 2RVP | NMR | DNA | null | 2rvp_CC_1_DC_8_2_DC_23 | stacked in. metal. | 0.116 | 10.391 |
| CT | 5CNQ | X-ray | Protein#DNA | 2.6 | 5cnq_CT_R_DC_4_H_DT_27 | non h-bonded WC conformation | 0.028 | 9.312 |
| CT | 5HP4 | X-ray | Protein#DNA | 1.86 | 5hp4_CT_X_DC_4_X_DT_7 | Constricted 2 hbond conf | 0.342 | 8.421 |
| CT | 5HP4 | X-ray | Protein#DNA | 1.86 | 5hp4_CT_X_DT_7_X_DC_4 | Constricted 2 hbond conf | -0.342 | 8.421 |
| CT | 5WSQ | X-ray | DNA | 1.05 | 5wsq_CT_A_DC_3_B_DT_6 | Stacked in WC. Mercury. | -1.907 | 9.864 |
| CT | 5WSQ | X-ray | DNA | 1.05 | 5wsq_CT_A_DT_6_B_DC_3 | Stacked in WC. Mercury. | 1.97 | 9.972 |
| CT | 1FKY | NMR | DNA | null | 1fky_CT_A_DC_6_B_DT_17 | Constricted 2 hbond & single h-bond | 0.252 | 8.882 |
| GG | 1D80 | X-ray | DNA | 2.2 | 1d80_GG_A_DG_4_B_DG_21 | G(syn).G(anti) | 0.717 | 10.737 |
| GG | 1D80 | X-ray | DNA | 2.2 | 1d80_GG_A_DG_9_B_DG_16 | G(syn).G(anti) | -0.868 | 11.23 |
| GG | 1OH7 | X-ray | Protein#DNA | 2.5 | 1oh7_GG_E_DG_9_F_DG_22 | Distorted G(syn).G(anti) | 2.303 | 13.209 |
| GG | 3DPG | X-ray | Protein#DNA | 1.91 | 3dpg_GG_C_DG_6_D_DG_13 | G(syn).G(anti) | -0.769 | 10.994 |
| GG | 3DPG | X-ray | Protein#DNA | 1.91 | 3dpg_GG_C_DG_13_D_DG_6 | G(syn).G(anti) | 1.549 | 11.122 |
| GG | 3LSP | X-ray | Protein#DNA | 2.66 | 3lsp_GG_B_DG_10_B_DG_21 | G(anti).G(anti) | -0.424 | 10.744 |
| GG | 3LSP | X-ray | Protein#DNA | 2.66 | 3lsp_GG_B_DG_21_B_DG_10 | G(anti).G(anti) Weird | 0.424 | 10.744 |
| GG | 4XZF | X-ray | Protein#DNA | 1.38 | 4xzf_GG_B_DG_7_B_DG_7 | G(syn).G(anti) & G(anti).G(syn) | 0 | 11.061 |
| GG | 5DBC | X-ray | Protein#DNA | 2.4 | 5dbc_GG_T_DG_13_P_DG_4 | G(syn).G(anti) | -1.702 | 10.847 |
| GT | 113D | X-ray | DNA | 2.5 | 113d_GT_A_DG_4_B_DT_21 | Wobble | -2.486 | 10.367 |
| GT | 113D | X-ray | DNA | 2.5 | 113d_GT_A_DT_9_B_DG_16 | Wobble | 2.717 | 10.274 |
| GT | 1D92 | X-ray | DNA | 2.25 | 1d92_GT_A_DG_3_B_DT_14 | Wobble | -2.581 | 10.451 |
| GT | 1D92 | X-ray | DNA | 2.25 | 1d92_GT_A_DT_6_B_DG_11 | Wobble | 2.138 | 10.331 |
| GT | 1E3M | X-ray | Protein#DNA | 2.2 | 1e3m_GT_E_DG_9_F_DT_22 | Distorted single h-bond conformation | 4.649 | 12.456 |
| GT | 1NG9 | X-ray | Protein#DNA | 2.6 | 1ng9_GT_E_DG_9_F_DT_22 | Distorted single h-bond conformation | 4.956 | 12.614 |
| GT | 1NK9 | X-ray | Protein#DNA | 1.9 | 1nk9_GT_B_DG_27_C_DT_6 | Inverse wobble | 1.906 | 10.922 |
| GT | 1NKB | X-ray | Protein#DNA | 2 | 1nkb_GT_B_DG_26_C_DT_7 | Inverse wobble | 2.082 | 11.203 |
| GT | 1NKC | X-ray | Protein#DNA | 1.8 | 1nkc_GT_B_DG_11_C_DT_32 | Wobble | -2.171 | 10.469 |
| GT | 1W7A | X-ray | Protein#DNA | 2.27 | 1w7a_GT_E_DG_9_F_DT_22 | Distorted single h-bond conformation | 4.78 | 12.415 |
| GT | 1WB9 | X-ray | Protein#DNA | 2.1 | 1wb9_GT_E_DG_9_F_DT_22 | Distorted single h-bond conformation | 4.854 | 12.566 |
| GT | 1WBB | X-ray | Protein#DNA | 2.5 | 1wbb_GT_E_DG_9_F_DT_22 | Distorted single h-bond conformation | 5.015 | 12.499 |
| GT | 1WBD | X-ray | Protein#DNA | 2.4 | 1wbd_GT_E_DG_9_F_DT_22 | Distorted single h-bond conformation | 4.932 | 12.653 |
| GT | 2O8B | X-ray | Protein#DNA | 2.75 | 2o8b_GT_E_DG_8_F_DT_23 | Distorted single h-bond conformation | 5.206 | 12.104 |
| GT | 2XM3 | X-ray | Protein#DNA | 2.3 | 2xm3_GT_K_DG_20_K_DT_33 | Single h-bond wobble with water | -2.661 | 9.746 |
| GT | 2XM3 | X-ray | Protein#DNA | 2.3 | 2xm3_GT_M_DG_20_M_DT_33 | Single h-bond wobble with water | -3.058 | 9.742 |
| GT | 2XMA | X-ray | Protein#DNA | 2.3 | 2xma_GT_C_DG_20_C_DT_33 | Single h-bond wobble with water | -2.392 | 9.751 |
| GT | 2XMA | X-ray | Protein#DNA | 2.3 | 2xma_GT_D_DG_20_D_DT_33 | Single h-bond wobble | -2.388 | 9.757 |
| GT | 2XMA | X-ray | Protein#DNA | 2.3 | 2xma_GT_G_DG_20_G_DT_33 | Single h-bond wobble | -2.884 | 9.765 |
| GT | 2XMA | X-ray | Protein#DNA | 2.3 | 2xma_GT_H_DG_20_H_DT_33 | Single h-bond wobble | -2.896 | 9.765 |
| GT | 2XO6 | X-ray | Protein#DNA | 1.9 | 2xo6_GT_B_DG_20_B_DT_33 | Single h-bond wobble with water | -2.705 | 9.818 |
| GT | 2XO6 | X-ray | Protein#DNA | 1.9 | 2xo6_GT_E_DG_20_E_DT_33 | Single h-bond wobble with water | -2.88 | 9.724 |
| GT | 2XQC | X-ray | Protein#DNA | 1.9 | 2xqc_GT_B_DG_20_B_DT_33 | Single h-bond wobble with water | -2.686 | 9.695 |
| GT | 2XQC | X-ray | Protein#DNA | 1.9 | 2xqc_GT_E_DG_20_E_DT_33 | Single h-bond wobble with water | -2.758 | 9.741 |
| GT | 3H25 | X-ray | Protein#DNA | 2.7 | 3h25_GT_C_DT_11_C_DG_23 | Wobble | 2.484 | 10.39 |
| GT | 3JXY | X-ray | Protein#DNA | 1.5 | 3jxy_GT_B_DG_7_C_DT_6 | Super weird distorted conformation | 7.505 | 11.212 |
| GT | 3K0S | X-ray | Protein#DNA | 2.2 | 3k0s_GT_E_DG_9_F_DT_22 | Distorted single h-bond conformation | 5.009 | 12.606 |
| GT | 3VXV | X-ray | Protein#DNA | 2 | 3vxv_GT_B_DG_10_C_DT_4 | Wobble | -2.325 | 10.647 |
| GT | 3VYQ | X-ray | Protein#DNA | 2.53 | 3vyq_GT_B_DG_7_C_DT_5 | Wobble | -2.206 | 10.717 |
| GT | 4FJ8 | X-ray | Protein#DNA | 2.19 | 4fj8_GT_T_DT_9_P_DG_111 | Inverse wobble | -2.031 | 11.242 |
| GT | 4M3U | X-ray | Protein#DNA | 2.07 | 4m3u_GT_T_DG_7_P_DT_113 | Wobble | -1.945 | 10.296 |
| GT | 4M3W | X-ray | Protein#DNA | 2.1 | 4m3w_GT_T_DG_8_P_DT_112 | Wobble | -2.109 | 10.312 |
| GT | 4M3X | X-ray | Protein#DNA | 2.2 | 4m3x_GT_T_DG_9_P_DT_111 | Wobble | -1.847 | 10.512 |
| GT | 4M41 | X-ray | Protein#DNA | 2.15 | 4m41_GT_T_DT_7_P_DG_113 | Tautomer | -0.091 | 10.745 |
| GT | 4M42 | X-ray | Protein#DNA | 2.04 | 4m42_GT_T_DT_8_P_DG_112 | Wobble | 2.145 | 10.575 |
| GT | 4M45 | X-ray | Protein#DNA | 1.89 | 4m45_GT_T_DT_9_P_DG_111 | Inverse wobble | -2.252 | 11.37 |

| | | | | | | | | |
|---|---|---|---|---|---|---|---|---|
| GT | 4OSQ | X-ray | Protein#DNA | 2.26 | 4osq_GT_I_DG_7_J_DT_-7 | Tautomer | -0.278 | 10.753 |
| GT | 4OSR | X-ray | Protein#DNA | 1.94 | 4osr_GT_I_DG_7_J_DT_-7 | Tautomer | 0.205 | 10.814 |
| GT | 5D9I | X-ray | Protein#DNA | 1.7 | 5d9i_GT_X_DT_14_Y_DG_9 | Wobble | 2.177 | 10.486 |
| GT | 5DBA | X-ray | Protein#DNA | 1.97 | 5dba_GT_T_DT_13_P_DG_4 | Wobble | 2.318 | 10.344 |
| GT | 1BJD | NMR | DNA | null | 1bjd_GT_A_DG_4_B_DT_9 | Wobble | -2.59 | 10.318 |
| GT | 1BJD | NMR | DNA | null | 1bjd_GT_A_DT_9_B_DG_4 | Wobble | 2.591 | 10.319 |
| GT | 1KKW | NMR | DNA | null | 1kkw_GT_A_DT_4_B_DG_7 | Bifurcated | 1.171 | 10.807 |
| GT | 1KKW | NMR | DNA | null | 1kkw_GT_A_DG_7_B_DT_4 | Bifurcated | -1.169 | 10.81 |
| TT | 1MNV | X-ray | Protein#DNA | 2.6 | 1mnv_TT_A_DT_5_B_DT_14 | Wobble | 2.252 | 8.365 |
| TT | 1P51 | X-ray | Protein#DNA | 2.5 | 1p51_TT_E_DT_11_F_DT_11 | Wobble | -2.357 | 8.268 |
| TT | 1P71 | X-ray | Protein#DNA | 1.9 | 1p71_TT_C_DT_11_D_DT_11 | Wobble | 2.264 | 8.413 |
| TT | 1P78 | X-ray | Protein#DNA | 2.25 | 1p78_TT_C_DT_11_D_DT_11 | Wobble | 2.117 | 8.599 |
| TT | 5GSK | X-ray | DNA | 1.05 | 5gsk_TT_A_DT_3_A_DT_6 | Stacked in WC. Mercury. | 0.081 | 10.235 |
| TT | 5GSK | X-ray | DNA | 1.05 | 5gsk_TT_A_DT_6_A_DT_3 | Stacked in WC. Mercury. | -0.081 | 10.235 |
| TT | 5WSP | X-ray | DNA | 1.5 | 5wsp_TT_A_DT_3_A_DT_6 | Wobble | 2.388 | 8.978 |
| TT | 5WSP | X-ray | DNA | 1.5 | 5wsp_TT_A_DT_6_A_DT_3 | Wobble | -2.388 | 8.978 |
| TT | 5WSR | X-ray | DNA | 1.5 | 5wsr_TT_A_DT_3_A_DT_6 | Stacked in wobble. Mercury. | 1.584 | 9.569 |
| TT | 5WSR | X-ray | DNA | 1.5 | 5wsr_TT_A_DT_6_A_DT_3 | Stacked in wobble. Mercury. | -1.584 | 9.569 |
| TT | 5WSS | X-ray | DNA | 1.45 | 5wss_TT_A_DT_3_A_DT_6 | Stacked in WC. Mercury. | -0.045 | 10.077 |
| TT | 5WSS | X-ray | DNA | 1.45 | 5wss_TT_A_DT_6_A_DT_3 | Stacked in WC. Mercury. | 0.045 | 10.077 |
| TT | 1DSD | NMR | Protein#DNA | null | 1dsd_TT_A_DT_3_B_DT_14 | Wobble | 2.898 | 8.807 |
| TT | 1DSD | NMR | Protein#DNA | null | 1dsd_TT_A_DT_6_B_DT_11 | Wobble | -2.51 | 8.727 |
| TT | 2LL9 | NMR | DNA | null | 2ll9_TT_A_DT_6_B_DT_17 | Wobble | -2.842 | 9.673 |

a. For “Macromolecular Type”: “DNA” refers to free DNA, “Protein#DNA” refers to protein-bound DNA, “Protein#DNA#RNA” refers to protein-DNA-RNA complexes.
b. “Base pair ID” refers to a_b_c_d_e_f_g_h, where a = PDB ID, b = mismatch type, c = chain ID 1, d = residue name 1, e = residue number 1, f = chain ID 2, g = residue name 2, h = residue number 2.

#### Table S4. Summary of available mismatch structures in free DNA, from structural survey of PDB data

| Mismatch | Method | Total number of occurrences [a] | Number of occurrences of non-metal mediated mismatches [b] | Number of sheared conformations [c] | Number of occurrences in constricted conformation [d] | Number of occurrences of Hoogsteen mismatches [e] |
|---|---|---|---|---|---|---|
| G-G | X-ray | 2 | 2 | 0 | 0 | 2 |
| | NMR | 0 | 0 | 0 | 0 | 0 |
| A-A | X-ray | 0 | 0 | 0 | 0 | 0 |
| | NMR | 0 | 0 | 0 | 0 | 0 |
| A-G | X-ray | 6 | 6 | 0 | 0 | 6 |
| | NMR | 1 | 1 | 0 | 0 | 0 |
| G-T | X-ray | 4 | 4 | 4 | 0 | 0 |
| | NMR | 4 | 4 | 4 | 0 | 0 |
| A-C | X-ray | 2 | 2 | 2 | 0 | 0 |
| | NMR | 3 | 3 | 3 | 0 | 0 |
| T-T | X-ray | 8 | 2 | 2 | 2 | 0 |
| | NMR | 1 | 1 | 1 | 0 | 0 |
| C-C | X-ray | 0 | 0 | 0 | 0 | 0 |
| | NMR | 1 | 0 | 0 | 0 | 0 |
| C-T | X-ray | 2 | 0 | 0 | 0 | 0 |
| | NMR | 1 | 1 | 0 | 1 | 0 |

a. We only consider mismatches within canonical helical context, which refers to a single mismatch embedded by at least two Watson-Crick (WC) base pairs on both sides. Modified base were also excluded.
b. “Non-metal mediated mismatch” refers to a mismatch without heavy metal inserted in between the two bases.
c. “Sheared conformation” is defined as base pair shear parameters > 1.0 Å or < -1.0 Å.
d. “Constricted conformation” is defined by C1′-C1′ distance < 9.5 Å.
e. “HG mismatches” (Hoogsteen mismatches) refers to structures where one of the purine bases is in *syn* conformation.

#### Table S5. Transcription factors and DNA sites tested by SaMBA

**a.**

| Transcription factor | Structural protein family (nomenclature according to[1,2]) | Selected TF binding site(s) (numbers in parentheses show the version of the SaMBA DNA library containing each site) | Universal PBM E-score for core 8-mer site (E>0.35 is specific) |
|---|---|---|---|
| Ets1 | ETS | CGGAAGTGTCGTAAACGTCGGA**TATCCGGT**TCTTCTGGGCGCTAAGGGAGCTGACGGAGA(v1)<br>TAAAGGGAAGACGGTTGTGGTA**GTTCCGGG**TATTAACCTGTGTGGCGCGGGGTCTGAGGC(v1)<br>ACACACATGAGATATCTGATGC**CTTCCGTG**TGGTTAGCCAGGTGTAGTTTGGTCGTAGTG(v2)<br>GAGATGGCGCCTGTTATCTTCTGCCCT**GTTCCGGT**ATTCAATATCTCTGGGCTTGCTCAG(v2)<br>CATCTGCCCTGCGG**TTTCCTTC**TGCCTGATAAGATTTCACCTGTCATCTTTTTGCCTCCC(v2) | 0.48<br>0.44<br>0.43<br>0.47<br>0.36 |
| Cbf1 | bHLH | GATTAATTTCTGCCCTTATTC**GCACGTGC**CATAACACCTGTACCTCTCACACTAAAGACC(v1) | 0.49 |
| Creb1 | bZIP | CCCGG**TGACGT**AAACGTCGGATATCCGGTTCTTCTGGGCGCTAAGGGAGCTGACGGAGAA(v2) | 0.49 |
| E2f1 | E2F | GAGA**TGGCGCCT**GTTATCTTCTGCCCTGTTCCGGTATTCAATATCTCTGGGCTTGCTCAG(v2) | 0.48 |
| Stat3 | STAT | TGATACTATAAAAG**TTCCTGGA**ATTTCCGGATACATCGCAAATGGGCGGTAGGCGAAAAA(v4) | 0.48 |
| Ap2a | AP-2 | CATCTG**CCCTGCGG**TTTCCTTCTGCCTGATAAGATTTCACCTGTCATCTTTTTGCCTCCC(v3) | 0.48 |
| Six6 | Homeodomain | TTTCTTTCTTATGTGGTTTT**TGATACTC**TGGGACAGAATGTTTACGTGGTAAGTGTGTTG(v3) | 0.47 |
| Atf1 | bZIP | GCGTAGTGCCACCTGGTGGCCACGTGCCA**ACGTCAC**CTTCTGTATAGTGGTGGAGTGACC(v3) | 0.43 |
| Crem | bZIP | CCCGG**TGACGTAA**ACGTCGGATATCCGGTTCTTCTGGGCGCTAAGGGAGCTGACGGAGAA(v3) | 0.49 |
| Elk1 | ETS | GAGATGGCGCCTGTTATCTTCTGCCCT**GTTCCGGT**ATTCAATATCTCTGGGCTTGCTCAG(v3) | 0.49 |
| Egr1 | EGR family C2H2 ZF | TGATGGTG**GCGTGGGC**GATAAGGAGGGGTAAGCGGTAAGGCTAAAGGAGGGGAAAGAGGC(v2) | 0.49 |
| Gabpa | ETS | TAAAGGGAAGACGGTTGTGGTA**GTTCCGGG**TATTAACCTGTGTGGCGCGGGGTCTGAGGC(v1) | 0.44 |
| Gata1 | GATA | CGGAAGTGTCGTAAACGT**CGGATATC**CGGTTCTTCTGGGCGCTAAGGGAGCTGACGGAGA(v1) | 0.43 |
| Max | bHLH | TGTCGGGTAAGGGAATGTAGA**TCACGTGA**ATGAAAATTCAAGTGTTTTTCGAAAAGTTTC(v1) | 0.49 |
| Myc | bHLH | TGTCGGGTAAGGGAATGTAGA**TCACGTGA**ATGAAAATTCAAGTGTTTTTCGAAAAGTTTC(v1)<br>GCGTAGTGCCACCTGGTGGC**CACGTG**CCAACGTCACCTTCTGTATAGTGGTGGAGTGACC(v2) | 0.47<br>0.49 |
| Mad | bHLH | CACAATTTTTACAGTAATGTAG**CACGCG**TAACTCCTAAACTTTGTCATAATGGTTGAAAT(v1) | 0.47 |
| Runx1 | RUNX | TTTCTTTCTTA**TGTGGTTT**TTGATACTCTGGGACAGAATGTTTACGTGGTAAGTGTGTTG(v2) | 0.49 |
| p53 | p53 | GCAGA**CATGCCCGGGCATG**CCTCGTGCTCCTGGGGTTTGTCATAATGGTTGAAATATGGC(v3) | Ref [3] |
| CTCF | C2H2 ZF | GGCAG**CGCCCTCTACTGGC**AGCTGCACGTGGTGGCCGCGCGCGCCGAGGCGGTGAGCGTG(v2) | Ref [4] |
| RelA | Rel | CACCTGG**GGAATTTCC**GGGAAAGTCACCTTCTGTATAGTGGTGGAGTGACCGCGTAGTGC(v4) | Ref [5] |
| TBP | TBP | TGATAC**TATAAAAG**TTCCTGGAATTTCCGGATACATCGCAAATGGGCGGTAGGCGAAAAA(v4) | 0.47 |

**b.**

| Transcription factor | Non-specific sequence | Designed originally for |
|---|---|---|
| Ets1 | TGATGGTGGCGTGGGCGATAAGGAGGGGTAAGCGGTAAGGCTAAAGGAGGGGAAAGAGGC<br>TCAAAACCGCAGGGGGGCGCTATGGCGCCATGTCACTGATAGTCGGGTGTCAATGGAAAC<br>GTGCCCGGCGCCCACGCGGGCGACACCTCCTCGTCGGTTATGAAGGGTGAAAAGGCACTC<br>GGTTTGTGGCCACCAGGTGGCACTAGGAATGTTCTGTGTTTTATGTCCCACCACCCAAGG | Gata1/Egr1 (v2 seq 3)<br>Runx/CTCF/E2f1 (v2 seq 10)<br>Myc/Egr1 (v2 seq 5)<br>CTCF (v2 seq 12) |
| Cbf1 | CGGAAGTGTCGTAAACGTCGGATATCCGGTTCTTCTGGGCGCTAAGGGAGCTGACGGAGA | Ets1 (v1 seq 1) |
| Creb1 | GGCAGCGCCCTCTACTGGCAGCTGCACGTGGTGGCCGCGCGCGCCGAGGCGGTGAGCGTG | CTCF/Myc (v2 seq 9) |
| E2f1 | CATCTGCCCTGCGGTTTCCTTCTGCCTGATAAGATTTCACCTGTCATCTTTTTGCCTCCC | Runx1 (v2 seq 7) |
| Stat3 | TTCATTCTTTTATATGAGTCATTCATTCATTCATACCACAGTCCATGCCATCACGAAAAA | TBP (v4 seq2) |
| Ap2a | ACACACATGAGATATCTGATGCCTTCCGTGTGGTTAGCCAGGTGTAGTTTGGTCGTAGTG | Elk1/Gata (v3 seq **1**) |
| Six6 | ACACACATGAGATATCTGATGCCTTCCGTGTGGTTAGCCAGGTGTAGTTTGGTCGTAGTG | Elk1/Gata (v3 seq **1**) |
| Atf1 | AACTTTGATATCATTGCACGTGTTTACTTATTTAGAGGTATCGTTGTTCGACGTTTTAGG | Six6/Myc (v3 seq 4) |
| Crem | GTGCCCGGCGCCCACGCGGGCGACACCTCCTCGTCGGTTATGAAGGGTGAAAAGGCACTC | Myc/Egr1 (v3 seq 5) |
| Elk1 | GCGTAGTGCCACCTGGTGGCCACGTGCCAACGTCACCTTCTGTATAGTGGTGGAGTGACC | CTCF/Myc/Creb(v3 seq 8) |
| Egr1 | GGCAGCGCCCTCTACTGGCAGCTGCACGTGGTGGCCGCGCGCGCCGAGGCGGTGAGCGTG | CTCF/Myc (v2 seq 9) |
| Gabpa | TGTCGGGTAAGGGAATGTAGATCACGTGAATGAAAATTCAAGTGTTTTTCGAAAAGTTTC | Cbf1 (v1 seq9) |
| Gata1 | GCTGCTGCGGTGCATACCCCACCGGATGTGACCCCGTGATGCGGTCTTGATTCGCTTGAC | Ets1 (v1 seq2) |
| Max | CAGCATAGGACTTCTTCCCTGAAGCAATTGATAACCCCGGTCTACGATGCGTGGTTCCAA | Ets1 control (v1 seq6) |
| Myc | GAGATGGCGCCTGTTATCTTCTGCCCTGTTCCGGTATTCAATATCTCTGGGCTTGCTCAG | E2f1/Gata1 (v2 seq2) |
| Mad | CAGCATAGGACTTCTTCCCTGAAGCAATTGATAACCCCGGTCTACGATGCGTGGTTCCAA | Ets1 control (v1 seq6) |
| Runx1 | TGATGGTGGCGTGGGCGATAAGGAGGGGTAAGCGGTAAGGCTAAAGGAGGGGAAAGAGGC | Gata1/Egr1(v2 seq 3) |
| p53 | ACACACATGAGATATCTGATGCCTTCCGTGTGGTTAGCCAGGTGTAGTTTGGTCGTAGTG | Gata/Elk1 (v3 seq 1) |
| CTCF | TTTCTTTCTTATGTGGTTTTTGATACTCTGGGACAGAATGTTTACGTGGTAAGTGTGTTG | Runx1/Six6 (v2 seq 6) |
| RelA | CTTCGCGCGCGCGCATGTCCGCGGACATGCACACGCTGTGGCTAAGTTTCGGATAATCGG | p53/Stat3 (v4 seq 5) |
| TBP | CACCTGGGGAATTTCCGGGAAAGTCACCTTCTGTATAGTGGTGGAGTGACCGCGTAGTGC | Stat3 (v4 seq 4) |

#### Table S6. X-ray and NMR structures of protein-DNA complexes selected for structural analyses of mismatches that increase TF binding

| Transcription factor | Number of protein-DNA structures available | PDB ID for selected structure | Method | Resolution (Å) | Sequence (underlined regions mark the core binding site sequences, which are identical between the PDB structures and the sites tested in SaMBA) |
|---|---|---|---|---|---|
| Creb1 | 3 | 5ZK1 | X-ray | 3.05 | CTTGGCTGACGTCAGCCAAG |
| CTCF | 12 | 5T00 | X-ray | 2.19 | TAGCGCCCCCTGCTGGC |
| Egr1 | 5 | 1P47 | X-ray | 2.2 | TGGCGTGGGCGGCGTGGGCGT |
| Elk1 | 1 | 1DUX | X-ray | 2.1 | TGACCGGAAGTGT |
| Ets1 | 17 | 2STT | NMR | N/A | TCGAGCCGGAAGTTCGA |
| Gata1 | 6 | 3VEK | X-ray | 2.63 | GTCTTATCAGATGGACTC |
| Max | 5 | 1AN2 | X-ray | 2.9 | GTGTAGGTCACGTGACCTACAC |
| Myc | 2 | 1NKP | X-ray | 1.8 | GAGTAGCACGTGCTACTC |
| p53 | 32 | 3KZ8 | X-ray | 1.91 | GGGCATGCCCGGGCATGCCC |
| RelA | 8 | 5U01 | X-ray | 2.5 | AGCGGAAATTCCCGGGAATTTCCGCT |
| Runx1 | 15 | 1HJB | X-ray | 3 | AAGATTTCCAAACTCTGTGGTTGCG |
| TBP | 30 | 1CDW | X-ray | 1.9 | CTGCTATAAAAGGCTG |

#### Table S7. Mismatches that increase TF binding affinity and exhibit geometries similar to distorted base pairs in TF-bound DNA

| Protein | Mismatched site (mutated base is shown in red) [a] | Mimicked distortion | PDB structure information (ID and base-pair) | Is the mutated base in contact? [b] | Does the rescue mutation have the same effect as the mismatch? [c] |
|---|---|---|---|---|---|
| CTCF | 5'-AGCG**T**CC-3'<br>3'-TCGC**G**GG-5' | Shear | PDBID: 5T00<br>B.DC6 C.DG12 | No | No |
| Ets1 | 5'-TTC**G**GG-3'<br>3'-AAG**G**CC-5' | C1'-C1' distance (expansion) | PDBID: 2STT<br>B.DC7 C.DG28 | Yes | No |
| Max | 5'-TC**G**CGTGA-3'<br>3'-AG**T**GCACT-5' | Shear | PDBID: 1AN2<br>A.DA10 B.DT13 | No | No |
| p53 | 5'-C**T**TGCCCGGGCATGCC-3'<br>3'-G**T**ACGGGCCCGTACGG-5' | C1'-C1' distance (constriction) | PDBID: 3KZ8<br>C.DA5 D.DT16 | No | No |
| p53 | 5'-C**C**TGCCCGGGCATGCC-3'<br>3'-G**T**ACGGGCCCGTACGG-5' | C1'-C1' distance (constriction) | PDBID: 3KZ8<br>C.DA5 D.DT16 | No | No |
| p53 | 5'-CATGCCCGGGC**T**TGCC-3'<br>3'-GTACGGGCCCG**T**ACGG-5' | C1'-C1' distance (constriction) | PDBID: 3KZ8<br>C.DA15 D.DT6 | No | No |
| p53 | 5'-CATGCCCGGGC**C**TGCC-3'<br>3'-GTACGGGCCCG**T**ACGG-5' | C1'-C1' distance (constriction) | PDBID: 3KZ8<br>C.DA15 D.DT6 | No | No |
| RelA | 5'-GGGAATTTCC**T**-3'<br>3'-CCCTTAAAGG**C**-5' | Stacking | PDBID: 5u01<br>B.DG12 C.DC105 | Yes | No |
| RelA | 5'-GGGAATTTCC**A**-3'<br>3'-CCCTTAAAGG**C**-5' | Stacking | PDBID: 5u01<br>B.DG12 C.DC105 | Yes | No |
| RelA | 5'-GGGAATTTCC**C**-3'<br>3'-CCCTTAAAGG**C**-5' | Stacking | PDBID: 5u01<br>B.DG12 C.DC105 | Yes | No |
| Runx1 | 5'-TG**C**GGTT-3'<br>3'-AC**A**CCAA-5' | Shear | PDBID: 1HJB<br>H.DA9 G.DT19 | No | No[d] |
| TBP | 5'-CTATAAA**T**G-3'<br>3'-GATATTT**T**C-5' | C1'-C1' distance (constriction) | PDBID: 1CDW<br>B.DA11 C.DT106 | No | No |
| TBP | 5'-CTATAAAA**C**-3'<br>3'-GATATTTT**C**-5' | Stacking | PDBID: 1CDW<br>B.DG12 C.DC105 | No | No |
| TBP | 5'-CTATAAAA**T**-3'<br>3'-GATATTTT**C**-5' | Stacking | PDBID: 1CDW<br>B.DG12 C.DC105 | No | No |
| TBP | 5'-CTATAAAA**A**-3'<br>3'-GATATTTT**C**-5' | Stacking | PDBID: 1CDW<br>B.DG12 C.DC105 | No | No |

a. Regions shown are the core binding site sequences, as underlined in **Table S6** (column 6), which are identical between the X-ray/NMR structures and the sequences tested in SaMBA.
b. The intermolecular hydrogen bonds (H-bond) between amino acids and DNA bases were detected by a geometrically based algorithm using X3DNA-DSSR 6 with a distance cutoff of less than 4.0 Å. We then manually inspected all intermolecular H-bonds to remove obvious false positive H-bonds such as donor-donor and acceptor-acceptor pairs.
c. To determine whether the rescue mutation (which restores Watson-Crick base-pairing) also increases TF binding levels, we used mutation binding data (**Fig S8**, Methods) and the DNA binding motifs of the TF proteins (**Fig. S6**).
d. For this particular Runx1 binding site, the rescue mutation also increases the protein binding level, although to a lesser extent than the mismatch. We verified that the C-A mismatch increases Runx1 binding levels even relative to the mutated, higher affinity Runx1 site 5'-TGCGGTT-3' (**Data File 1**).
